# Functional memory T cells are derived from exhausted clones and expanded by checkpoint blockade

**DOI:** 10.1101/2025.02.10.637523

**Authors:** Colin J. Raposo, Patrick K. Yan, Andy Y. Chen, Saba Majidi, Kamir J. Hiam-Galvez, Ansuman T. Satpathy

## Abstract

Immune checkpoint blockade can facilitate tumor clearance by T cells, resulting in long term patient survival. However, the capacity of exhausted CD8^+^ T cells (Tex), present during chronic antigen exposure, to form memory after antigen clearance remains unclear. Here, we performed longitudinal single cell RNA/T cell receptor sequencing and ATAC-sequencing on antigen-specific T cells after the clearance of chronic lymphocytic choriomeningitis virus (LCMV) infection. These data revealed the formation of a robust population of memory CD8^+^ T cells that transcriptionally, epigenetically, and functionally resemble central memory T cells (Tcm) that form after clearance of acute infection. To lineage trace the origin and memory recall response of Tex-derived memory clones, we utilized T cell receptor sequencing over the course of primary infection and rechallenge. We show that chronic Tcm are a clonally distinct lineage of Tex derived from progenitor exhausted cells, persist long-term in the absence of antigen, and undergo rapid clonal expansion during rechallenge. Finally, we demonstrate that αPD-L1 immune checkpoint blockade after chronic LCMV infection preferentially expands clones which form Tcm after clearance. Together, these data support the concept that chronically stimulated T cells form *bona fide* functional memory T cells through an analogous differentiation pathway to acutely stimulated T cells, which may have significant implications for enhancing immune memory to cancer through checkpoint blockade and vaccination.

## Main

Persistent antigen present during cancer and chronic infection drives CD8^+^ T cell exhaustion – a cell state categorized by the expression of co-inhibitory receptors (e.g. PD-1, Tim-3, Lag-3) and decreased cytokine production, proliferation, and target cell killing^1–8^. Recent developments in high-throughput genomic technologies, including single cell RNA, T cell receptor (TCR), and assay for transposase-accessible chromatin (ATAC) sequencing have provided new insights into the functional subsets, molecular regulation, and clonal differentiation pathways of exhausted T cells (Tex). These studies have revealed that Tex are a functionally diverse cellular pool that includes hypofunctional terminally exhausted cells (Tex-Term), effector-like intermediate cells (Tex-Int), killer cell lectin-like receptor-expressing exhausted cells (Tex-KLR), and stem-like progenitor exhausted cells (Tex-Prog) that reseed other Tex populations and respond to immune checkpoint blockade (ICB; ref^9–16^). The ability to track individual T cell clones using TCR sequencing has provided even greater resolution of Tex differentiation and revealed that individual Tex clones can exhibit a bias towards either Tex-Term or Tex-KLR terminally differentiated states^11^. Finally, epigenetic profiling has demonstrated that Tex represent a distinct differentiation trajectory that is accompanied by genome-wide changes in chromatin accessibility that limit subsequent activation following antigen recognition^17,18^.

Despite these advances in understanding of Tex differentiation, the field lacks a similar level of insight into the long-term impact of chronic stimulation on the generation of T cell memory. After the clearance of antigen from acute infections or vaccination, the antigen-specific T cell compartment contracts, and the remaining cells form long-lived memory T cells^19,20^. Upon reexposure to antigen, memory T cells expand and differentiate more rapidly than naive T cells and provide protection against reinfection^21,22^. However, after the clearance of chronic antigens, antigen-specific T cells persist long-term, but are not thought to provide the enhanced protection after reinfection characteristic of memory T cells^23,24^. While persisting T cells can express some markers of T cell memory (e.g. CD127, CD62L, TCF-1) after chronic antigen clearance, their ability to re-expand is lower than acute memory T cells, and this functional difference has been explained by epigenetic differences resulting from chronic stimulation^4,23,25–28^. Namely, chronic memory-like cells have increased accessibility of loci which encode for exhaustion related transcription factors (e.g. *Tox, Tox2,* and *Batf*) and co-inhibitory receptors (e.g. *Pdcd1*, *Lag3*, and *Tigit*) and decreased accessibility of effector function-associated gene loci^25,28^. These changes reinforce a Tex cell state upon stimulation and lead to decreased expansion, effector cell differentiation, and production of proinflammatory cytokines, including interferon (IFN)-γ and tumor necrosis factor (TNF)-α^26–28^. However, to date, these functional and epigenetic studies have largely been performed on aggregate Tex populations, typically for a single TCR specificity, rather than on individual clones within a polyclonal T cell response. Thus, it is unclear whether all antigen-specific clones maintain a dysfunctional phenotype following clearance of chronic antigen or whether a subset of Tex clones may be able to generate functional memory T cells.

Here, we leveraged high-throughput single cell RNA and TCR sequencing (scRNA/TCR-seq) to understand the cellular and clonal heterogeneity of antigen-specific T cells after the clearance of chronic viral infection. We show that antigen-specific T cells form a population of central memory T cells (Tcm) that have a shared transcriptional and epigenomic profile with Tcm formed after the clearance of acute infection. Through longitudinal profiling of the TCR repertoire, we show that chronic Tcm (cTcm) derive from a distinct differentiation trajectory of Tex-Prog biased clones present during chronic infection, and upon rechallenge exhibit enhanced capacity to expand and differentiate, promoting viral clearance. Finally, utilizing longitudinal TCR repertoire analysis over the course of infection, we demonstrate that ICB preferentially expands and differentiates clones that form cTcm while preserving their memory function. Altogether, these findings demonstrate the capacity of a clonally-restricted subset of Tex clones to form highly functional memory cells and highlight the potential of therapeutic interventions to enhance memory responses to chronic infections or cancer.

### Antigen-specific memory T cells are formed after the clearance of chronic viral infection

To study the clonal differentiation patterns of antigen-specific T cells after the clearance of chronic antigen exposure, we performed scRNA/TCR-seq on antigen-specific T cells from the spleen after the peripheral clearance of chronic lymphocytic choriomeningitis virus clone-13 (LCMV-Cl13), achieved by 100 days post infection (dpi**; Extended Data Fig. 1a**; ref^1^). We analyzed transcriptomes of GP33:H2-D^b^ tetramer-reactive T cells (GP33^+^), GP33^+^ CD62L^+^ T cells (to enrich for memory-like cells), or GP33^+^ CD62L^-^ CD8^+^ T cells: (1) after peripheral clearance of LCMV-Cl13 (100+ dpi), (2) during LCMV-Cl13 infection (21-26 dpi), or (3) after clearance of the acute LCMV-Armstrong (LCMV-Arm; 60+ dpi; **Fig. 1a,b**, **Extended Data Fig. b-d**). Clustering of cells (n=34,269) identified distinct exhausted and memory T cell subsets based on differential gene expression (LogFC>0.25; **Fig. 1b,c**; **Supplementary Table 1**). During chronic infection, GP33^+^ T cells comprised Tex clusters, including Tex-Term, characterized by high expression of *Pdcd1* (encoding PD-1), *Tox,* and *Cd101* (**Fig. 1c,d**). Effector gene expression (*Cx3cr1, Zeb2, and Klrg1*) along with *Pdcd1* and *Tox* expression identified Tex-Int and Tex-KLR (**Fig. 1c,d**). Tex-Prog were identified by high expression of *Slamf6* (encoding Ly108), *Tcf7* (encoding T Cell Factor [TCF]-1), *Pdcd1*, and *Tox* (**Fig. 1c,d**). After LCMV-Arm clearance, GP33^+^ cells comprised canonical memory cell clusters, including Tcm (expressing *Sell* [encoding CD62L]*, Tcf7,* and *Il7r* [encoding CD127]) and effector memory cell (Tem; expressing *Il7r, Cx3cr1,* and *Zeb2*) states (**Fig. 1c,d**). Unexpectedly, while canonical Tex subsets found in earlier time points persisted post peripheral LCMV-Cl13 clearance, the majority of GP33^+^ cells clustered with either acute Tem or Tcm, suggesting that Tex can differentiate to form cells with transcriptional phenotypes that are highly similar to Tcm and Tem (**Fig. 1d**).

**Figure 1:**
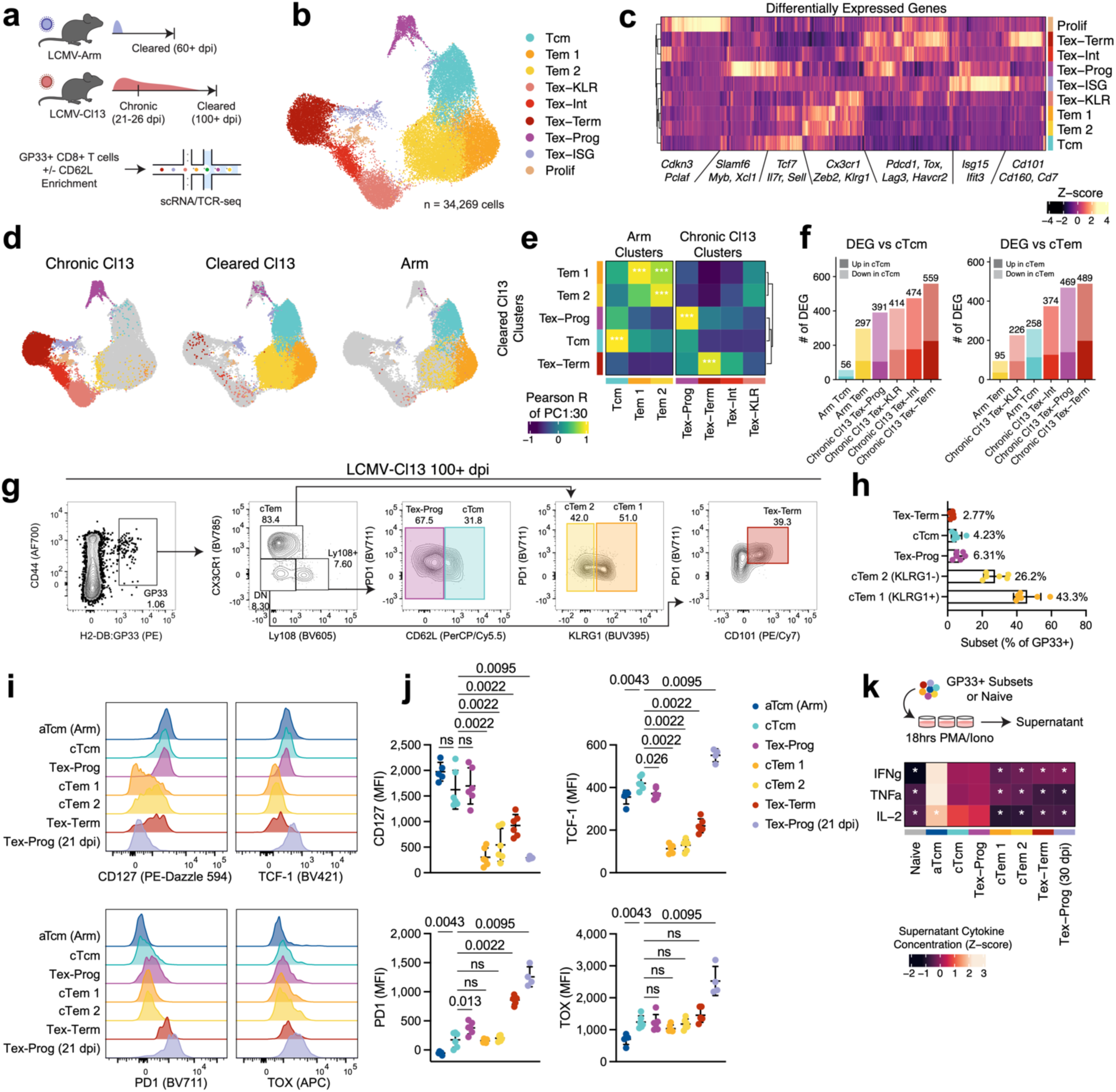
Tcm arise after the clearance of chronic viral infection. (a-f) scRNA/TCR-seq of GP33^+^, GP33^+^ CD62L^+^, and GP33^+^ CD62L^-^ CD8^+^ T cells sorted from the spleen at various timepoints after LCMV-Cl13 or LCMV-Arm. (a) Experimental schematic: LCMV-Arm and LCMV-Cl13 viral burden are graphically illustrated as shaded regions. (b) UMAP of all scRNA-seq cell profiles colored by cluster. (c) Differentially expressed genes per cell cluster. (d) UMAP of cells from each group colored by cluster. (e) Pearson correlation of PCs 1-30 between cleared LCMV-Cl13 clusters and chronic LCMV-Cl13 or LCMV-Arm clusters (Bonferonni corrected *P* values). (f) Number of differentially expressed genes (DEGs) between cTcm or cTem and cell subsets from chronic LCMV-Cl13 or LCMV-Arm. (a-f) n=8 cleared LCMV-Cl13, n=5 LCMV-Arm, n=8 chronic LCMV-Cl13. Data are pooled from two experiments. (g) Flow cytometry gating scheme for T cell subsets 100+ dpi with LCMV-Cl13 (gated on live CD8^+^). (h) Cell subset frequency of GP33^+^ cells 100+ dpi LCMV-Cl13. (i,j) CD127, TCF-1, PD-1, and TOX expression among GP33^+^ cell subsets from 100+ dpi LCMV-Cl13 or controls (aTcm LCMV-Arm 100+ dpi; Tex-Prog LCMV-Cl13 21 dpi). (g-j) Data shown as mean +/− SD. n=6 LCMV-Cl13 −100+ dpi, n=5 LCMV-Arm, n=4 LCMV-Cl13 21 dpi. Representative experiment of at least two independent experiments. *P* values calculated by Mann-Whitney U Test. (k) *In vitro* cytokine production by GP33^+^ subsets following 18 hour stimulation with PMA/Ionomycin. n=3-5 replicates per group. Cells pooled from multiple mice: n=11 LCMV-Cl13 −100+ dpi, n=4 LCMV-Arm 100+ dpi, n=6 LCMV-Cl13 30 dpi. Significance for each cytokine are shown versus cTcm, calculated by DESeq2. (a-k)*=*P*<0.05, **=*P*<0.01, ***=*P*<0.001, ns=*P*>0.05.

To better understand the transcriptional state of chronic memory cells, we compared cTcm and chronic Tem (cTem; defined by similarity to Tem in acute infection) to other subsets based on scRNA-seq profiles. Global comparisons by principal component analysis (PCA) showed significantly correlated gene expression between Tcm derived from both infection contexts (*P*<0.001; **Fig. 1e**). Likewise, gene expression profiles of cTem from both infection contexts were also significantly correlated (*P*<0.001; **Fig. 1e**). Next, we performed differential expression analysis between cTcm and acute Tcm (aTcm), as well as cTem and acute Tem (aTem; **Fig. 1f**). Overall, there were fewer differentially expressed genes (DEGs; *P*<0.01, Log_2_FC>0.25; **Supplementary Table 2**) between cTcm and aTcm (n=56) than between cTcm and every other Tex subset from chronic infection (n≥391; **Fig 1f**). Similarly, there were fewer DEGs between cTem and aTem (n=95) than between cTem and other Tex subsets (n≥226; **Fig 1f**). The few DEGs that were more highly expressed in chronic memory cells were consistent with Tex origin, including *Tox*, *Pdcd1*, and the interferon stimulated gene *Ifi27l2a*, which were higher in cTcm/cTem versus aTcm/aTem (**Supplementary Table 2**). Conversely, aTcm and aTem had increased expression of activator protein-1 (AP-1) family transcription factors (*Jun, Junb, Jund*) and the chromatin remodeler *Satb1*, compared to cTcm and cTem, consistent with increased AP-1 activity in effector cells compared to Tex (**Supplementary Table 2**; ref^29^). Finally, we compared expression of a module of core exhaustion genes (including *Tox, Pdcd1, Tigit,* and *Lag3*; **Supplementary Table 7**; ref^11^) between cells subsets post-LCMV-Cl13 clearance, which showed that cTcm cells had a lower exhaustion module score (−0.077) than Tex clusters (Tex-Term=0.84, Tex-Prog=0.081, Tex-Int=0.39, Tex-KLR=0.076), and the exhaustion module score of cTcm was similar to that of aTcm (−0.12; **Extended Data Fig. 1e**).

To validate the presence of chronic cTcm using an orthogonal assay, we performed flow cytometry on GP33^+^ T cells from the same time points as scRNA/TCR-seq. Consistent with the scRNA/TCR-seq data, GP33^+^ T cells had significantly decreased PD-1^+^ cell frequency (FC=3.34 *P*=0.0095) and increased CD127^+^ cell frequency (FC=8.58 *P*=0.0095) at 100+ dpi compared to 21 dpi (**Extended Data Fig. 2a**). Similarly, we observed a significant increase in the frequency of CD127^+^ CD62L^+^ T cells (canonical Tcm markers) from 21 to 100+ dpi (FC=4.01 *P*=0.0095; **Extended Data Fig. 2a**). CD127^+^ CD62L^+^ T cells comprised 6.57% of GP33^+^ T cells (**Extended Data Fig. 2a**). We assessed the frequency of CD127^+^ CD62L^+^ T cells at later time points post-peripheral LCMV clearance and observed that they persisted for at least 350 dpi, suggesting long-term durability of cTcm, similar to Tcm formed after acute infection (**Extended Data Fig. 2b**; ref^30^).

To further evaluate functional differences and validate DEGs of chronic memory cells, we developed a flow cytometry gating scheme to identify and compare GP33^+^ cTcm, cTem, Tex-Prog, and Tex-Term at 100+ dpi to aTcm from LCMV-Arm (GP33^+^ CD62L^+^ CXCR3^+^) and Tex-Prog from ongoing LCMV-Cl13 infection (GP33^+^ CX3CR1^-^ Ly108^+^; **Fig. 1g**). Consistent with scRNA/TCR-seq, GP33^+^ cells were predominantly cTem at 100+ dpi (68.45%), with cTcm (4.23%), Tex-Prog (6.31%), and Tex-Term (2.78%) present at lower relative frequencies (**Fig. 1h**). TCF-1 expression was significantly higher in cTcm than other subsets from cleared LCMV-Cl13 (Tex-Prog: FC=1.13 *P=*0.026, Tex-Term: FC=1.9 *P=*0.0022, Tem 1: FC=3.71 *P=*0.0022, Tem 2: FC=3.25 *P=*0.0022; **Fig. 1i,j**). cTcm had significantly lower TOX (FC=2.37 *P*=0.0095) and PD-1 (FC=8.38 *P*=0.0095) expression relative to Tex-Prog from 21 dpi. Consistent with scRNA/TCR-seq data, cTcm had elevated PD-1 expression (*P*=0.043) compared to aTcm but decreased expression compared to Tex subsets (Tex-Prog: FC=2.14 *P*=0.013, Tex-Term: FC=4.9 *P*=0.0022, Tex-Prog 21 dpi: FC=8.38 *P*=0.0095; **Fig. 1i,j**). Finally, we evaluated the function of GP33^+^ cell subsets *in vitro* by measuring cytokine secretion following stimulation with PMA/Ionomycin. cTcm secreted similar amounts of IFNγ and TNFα as aTcm (*P*>0.05) and significantly higher levels of IFNγ, Interleukin (IL)-2, and TNFα compared to cTem or Tex-Prog from ongoing LCMV-Cl13 infection (*P*<0.05; **Fig. 1k**). Altogether, these data suggest that cTcm formed after the clearance of chronic LCMV infection are transcriptionally, phenotypically, and functionally similar to aTcm.

### cTcm are derived from Tex-Prog biased clones

Memory and exhausted T cells achieve population-level functional diversity through the differentiation of individual clones with distinct fate distributions^11,31–33^. To assess behaviors of individual T cell clones after peripheral LCMV-Cl13 clearance, we analyzed the scRNA/TCR-seq data and identified 246 expanded *Tcra/Tcrb* clones in scRNA/TCR-seq (n>1 cell per clone, average 35 clones per mouse). To identify shared clonal differentiation patterns, we clustered clones based on intraclonal phenotype proportions (**Fig. 2a, Extended Data Fig. 3a**). This analysis revealed five distinct clonal differentiation patterns of GP33^+^ T cells after chronic antigen clearance: cTcm biased clones (n=66), Tem 1 biased clones (n=49), Tem 2 biased clones (n=75), Tex biased clones (n=7), and divergent clones with no preferential fate bias (n=49; **Fig. 2a, Extended Data Fig. 3a**). We next performed bulk TCR β-chain sequencing (TCR-seq) on sorted GP33^+^ T cell subsets (cTcm, Tem 1, Tem 2, Tex-Prog, Tex-Term; **Extended Data Fig. 3b**) for greater TCR sequencing depth and clonal resolution (**Fig. 2b-d**). In this dataset, we identified 190 expanded clones (n>4 cells per clone, average 63 clones per mouse; **Fig. 2b**). Clustering of expanded clones from bulk TCR-seq further confirmed the presence of cTcm biased (n=25), Tem 1 biased (n=41), Tem 2 biased (n=39), and divergent clones (n=48; **Fig. 2b**). In addition, the depth of this dataset also enabled the identification of Tex-Prog biased (n=9) and Tex-Term biased (n=28) biased clones, which were not observed in scRNA/TCR-seq due to low sampling of small clones (**Fig. 2b**). Across both scRNA/TCR-seq and bulk TCR-seq data, the majority of cells in cTcm biased clones were cTcm (median=86.6% cTcm in scRNA/TCR-seq, median=66.5% in bulk TCR-seq; **Fig. 2b**, **Extended Data Fig. 3a**). To quantitatively assess intraclonal phenotype distributions, we asked whether clones in each cluster were enriched for a dominant phenotype compared to a background distribution (**Fig. 2c**). Indeed, we found that the majority of clones in Tex-Prog, cTcm, Tex-Term, and Tem 2 biased clusters were significantly enriched for their respective cell subsets (*P*<0.05 hypergeometric test), while only 10.4% of divergent clones showed significant enrichment (**Fig. 2c**). Finally, the cTcm subset had low clonal overlap with other T cell subsets, as measured by Morisita-Horn index (Median=0.66; **Fig. 2d**). In contrast, there was high clonal overlap between Tex-Prog / Tex-Term (0.91) and cTem 1 / cTem 2 subsets (0.94; **Fig. 2d**). These results demonstrate that cTcm represent a distinct clonal Tex differentiation trajectory and suggest the presence of clonally restricted cTcm precursors during chronic infection.

**Figure 2:**
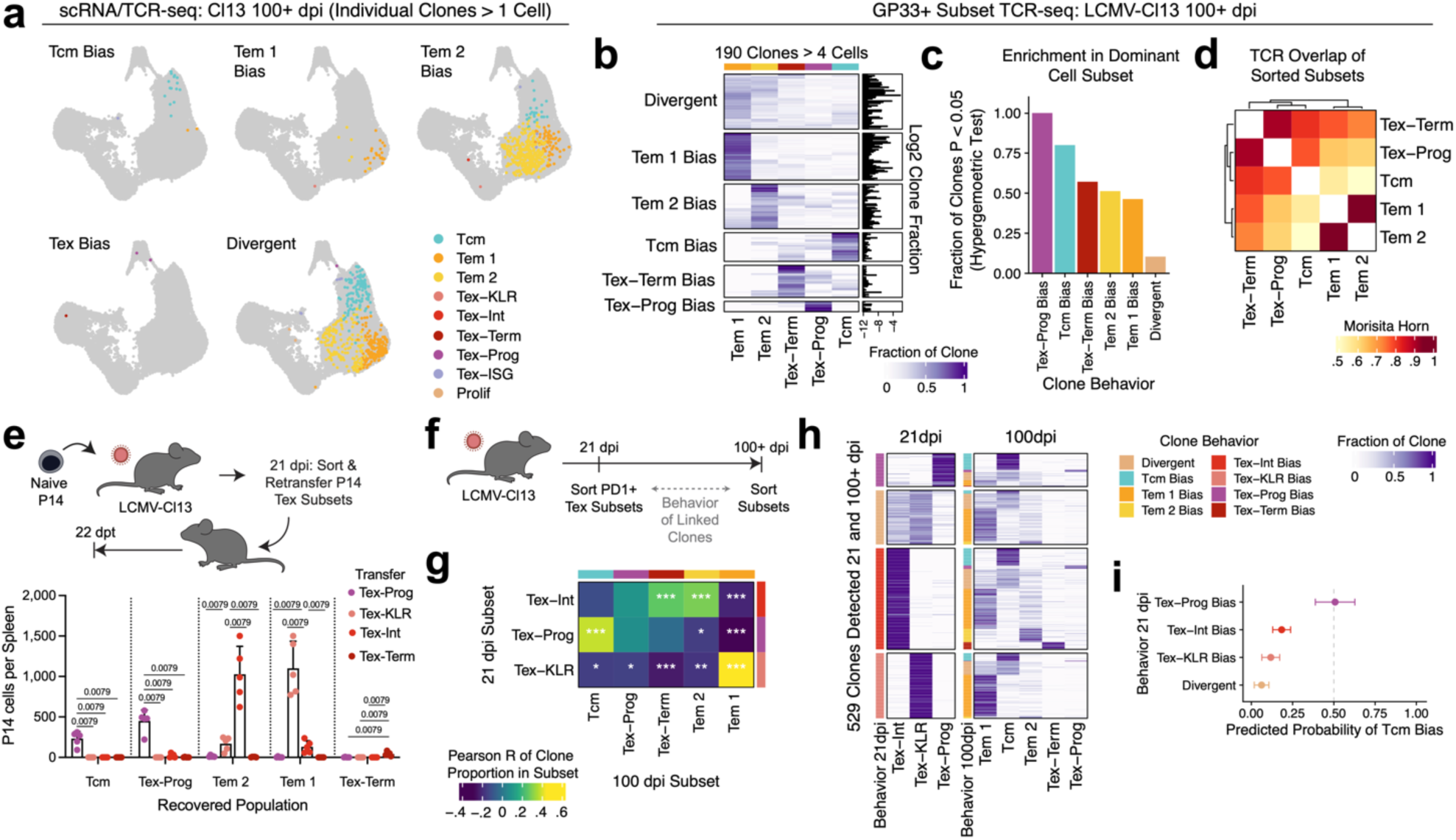
Chronic Tcm originate from Tex-Prog biased clones. (a) Representative clones of GP33^+^ CD8^+^ T cells from scRNA/TCR-seq. Clones are labelled based on k-means clustering of phenotype proportions among all clones from LCMV-Cl13 100+ dpi. See **Extended Data Fig. 3a**. (b-d) Bulk TCR-seq of GP33^+^ subsets isolated 100+ dpi with LCMV-Cl13 (n=3 mice). (b) Phenotype distribution of individual TCR clones across subsets, grouped by clonal behavior determined by k-means clustering. (c) Fraction of clones with statistically significant enrichment (*P*<0.05 Hoechberg corrected hypergeometric test) for the clone’s dominant cell subset. (d) Morisita-Horn TCR overlap index of sorted subsets from bulk TCR-seq. (e) Subsets of exhausted P14 were isolated from LCMV-Cl13 infected mice 21 dpi and 4.5×10^4^ cells were transferred into infection free secondary hosts. Flow cytometry quantification of P14 of cell subsets was performed 22 days post transfer (dpt). Bars show mean + standard deviation. n=5 mice per group. Representative experiment of two independent experiments. *P* values calculated by Mann-Whitney U Test. (f-i) T cell clones from LCMV-Cl13 infected mice were subset sorted from the blood 21 dpi and spleen 100+ dpi, and clonal behaviors were analyzed across timepoints by TCR-seq. (g) Pearson correlation of clone frequency in each subset between timepoints (Bonferonni corrected *P* values). (h) Phenotype distribution of individual TCR clones grouped by clonal behavior at 28 dpi and 100+ dpi. Color bars indicate clone behavior at each timepoint. (i) Probability of clones within different exhausted clone behaviors transitioning to cTcm bias after peripheral viral clearance. Probability +/− 95% CI calculated by logistic regression. (f-i) n=8 mice. (a-i)*=*P*<0.05, **=*P*<0.01, ***=*P*<0.001, ns=*P*>0.05.

To confirm that the cTcm biased clones were derived from Tex, rather than newly primed after the chronic phase of infection, we compared the TCR repertoire of GP33^+^ CD8^+^ T cells from peripheral blood 28 dpi to the TCR repertoire in the spleen from the same mice 100+ dpi (**Extended Data Fig. 3c,d**). We detected 25 cTcm biased clones at 100+ dpi, 14 of which were detected in the blood at 28 dpi (56%, 95% confidence interval [CI] = +/−19.5% by logistic regression), confirming that cTcm can indeed emerge from the exhausted T cell pool (**Extended Data Fig. 3d**). To determine whether a specific Tex subset preferentially forms cTcm, we transferred antigen-specific P14 Tex-Prog, Tex-Int, Tex-KLR, and Tex-Term from 21 dpi of LCMV-Cl13 infection into infection-free mice (**Fig. 2e**, **Extended Data Fig. 4e-g**). 22 days post transfer, we isolated the progeny of transferred cells and found that cTcm were derived exclusively from Tex-Prog (median=30.9% of Tex-Prog derived cells vs. 0% for other subsets; *P*=0.0079; **Fig. 2e**, **Extended Data Fig. 4f,g**)

Since not all transferred Tex-Prog formed CD62L^+^ cTcm, we asked whether a subset of clones that generate Tex-Prog may preferentially form cTcm after viral clearance. In murine cytomegalovirus infection, T cell clones that express memory associated genes early during the T cell response are enriched in the memory pool, so we hypothesized that clones with Tex-Prog bias may follow a parallel differentiation trajectory to form cTcm after chronic infection^34^. To test this, we first analyzed scRNA/TCR-seq data to determine if there were Tex-Prog biased clones during chronic infection, similar to the Tex-Prog and cTcm biased clones observed at later time points after clearance. By clustering clones based on intraclonal cluster proportion, we found a subset of clones (n=32) that were ≥50% Tex-Prog (**Extended Data Fig. 4a,b**). Next, to test if Tex-Prog biased clones preferentially formed cTcm, we longitudinally traced clones from the chronic phase of infection to after peripheral viral clearance in the same mice. We sorted PD-1^+^ Tex subsets (Tex-Int, Tex-KLR, and Tex-Prog) from the blood at 21 dpi, and after peripheral antigen clearance at 100 dpi, we performed TCR-seq of T cell subsets (cTcm, cTem 1, cTem 2, Tex-Prog, and Tex-Term) and compared clonal behaviors between timepoints (**Fig. 2f**, **Extended Data Fig. 4c**). We analyzed PD-1^+^ cells, rather than GP33^+^ cells, to evaluate T cells with a wider range of TCR avidity for antigen, instead of only high avidity clones that bind tetramer^35^. To validate this system prior to performing longitudinal clone tracing, we first analyzed TCR-seq data from PD-1^+^ T cell subsets in the spleen and blood of the same mouse at 28 dpi, which demonstrated that clone size and Tex-Prog, Tex-Int, and Tex-KLR phenotype distribution were significantly correlated between tissues (*P*<0.0001), suggesting that blood sampling accurately captures clonal behaviors of splenic Tex at a given time point (**Extended Data Fig. 4c-h**). Of note, and as previously described, there were few Tex-Term present in the peripheral blood, and we observed that splenic Tex-Term frequency showed the highest correlation with Tex-Int frequency in the blood (**Extended Data Fig. 4g,h**; ref^36^).

In the longitudinal data, we first replicated the results of the adoptive cell transfer experiments by analyzing clonal relationships between cell subsets at different timepoints. We calculated the subset frequency of each clone at both timepoints and then assessed the correlations of each subset frequency at 21 and 100+ dpi. Strikingly, intraclonal frequency of the Tex-Prog subset at 21 dpi significantly correlated with a clone’s cTcm frequency at 100+ dpi (R=0.39 *P*<0.001), supporting the link between the Tex-Prog and cTcm (**Fig. 2g**). In contrast, the intraclonal frequency of Tex-KLR from 21 dpi was significantly correlated with cTem 1 frequency after viral clearance (R=0.43 *P*<0.001), and Tex-Int frequency at 21 dpi was significantly correlated with cTem 2 (R=0.28 *P*<0.001) and Tex-Term (R=0.25 *P*<0.001) frequency after viral clearance (**Fig. 2g**). Finally, we examined the differentiation patterns of individual clones between time points. We isolated clones that were detected at both timepoints (n=529) and clustered clones based on intraclonal phenotype proportion from the blood at 21 dpi to identify Tex-Prog biased, Tex-KLR biased, Tex-Int biased, and divergent clonal behaviors (**Fig. 2h**). Similarly, we clustered the longitudinally traced clones based on phenotype proportion at 100+ dpi and identified cTcm biased, cTem 1 biased, cTem 2 biased, Tex-Prog biased, Tex-Term biased, and divergent clones (**Fig. 2h**). To identify the origin of cTcm biased clones, we compared the behavior of each clone at both timepoints and found that approximately 50% of Tex-Prog biased clones (34/69 clones; 50.7% +/−11.9% CI by logistic regression) detected in the blood at 21 dpi became cTcm biased clones after peripheral viral clearance at 100+ dpi (**Fig. 2i**). Conversely, divergent clones detected in the blood, many of which contain Tex-Prog, rarely formed cTcm biased clones (7/113 divergent clones, 5/32 Tex-Prog containing divergent clones; **Fig. 2h**). Similarly, only a small fraction of Tex-Int (36/211 clones) or Tex-KLR (16/136 clones) formed cTcm biased clones after viral clearance (**Fig. 2h**). These findings demonstrate that a subset of Tex clones with Tex-Prog bias preferentially gives rise to clonally restricted cTcm, highlighting the role of clonal diversification in the generation of memory cells after clearance of chronic antigen.

### Shared epigenomic cell states of cTcm and aTcm

Since chronic antigen stimulation can lead to changes in Tex chromatin state that may not be immediately reflected in RNA expression, we asked whether these changes were also observed in cTcm^25,28^. We performed epigenomic profiling of antigen-specific T cell subsets using ATAC-seq to measure the accessibility of *cis*-regulatory elements associated with exhaustion^37^. We sorted GP33^+^ CD8^+^ T cell subsets from the spleen after peripheral LCMV-Cl13 clearance, as well as GP33^+^ Tex populations from LCMV-Cl13 infected mice 28 dpi and GP33^+^ memory T cells after LCMV Arm infection for comparison (**Fig. 3**, **Extended Data Fig. 5a**). We first analyzed the 10,000 most variable open chromatin regions (OCRs; 500bp non-overlapping genomic regions) between samples and performed PCA and clustering, which identified five clusters of samples with global similarities in chromatin accessibility (**Fig. 3a**). Strikingly, cTcm samples clustered exclusively with aTcm, rather than with Tex-Prog from chronic infection, including CD62L^+^ Tex-Prog, which have recently been identified as a more functional population of Tex-Prog (**Fig. 3a**; ref^16^). Similarly, cTem samples clustered with aTem, reinforcing the notion that Tex can differentiate into Tem/Tcm in a parallel manner to acutely stimulated cells after chronic antigen clearance. Tex-Prog that persisted past chronic antigen clearance clustered on their own, and PCA analysis showed that this subset exhibits an intermediate state between Tex-Prog from active infection and Tcm (**Fig. 3a**).

**Figure 3:**
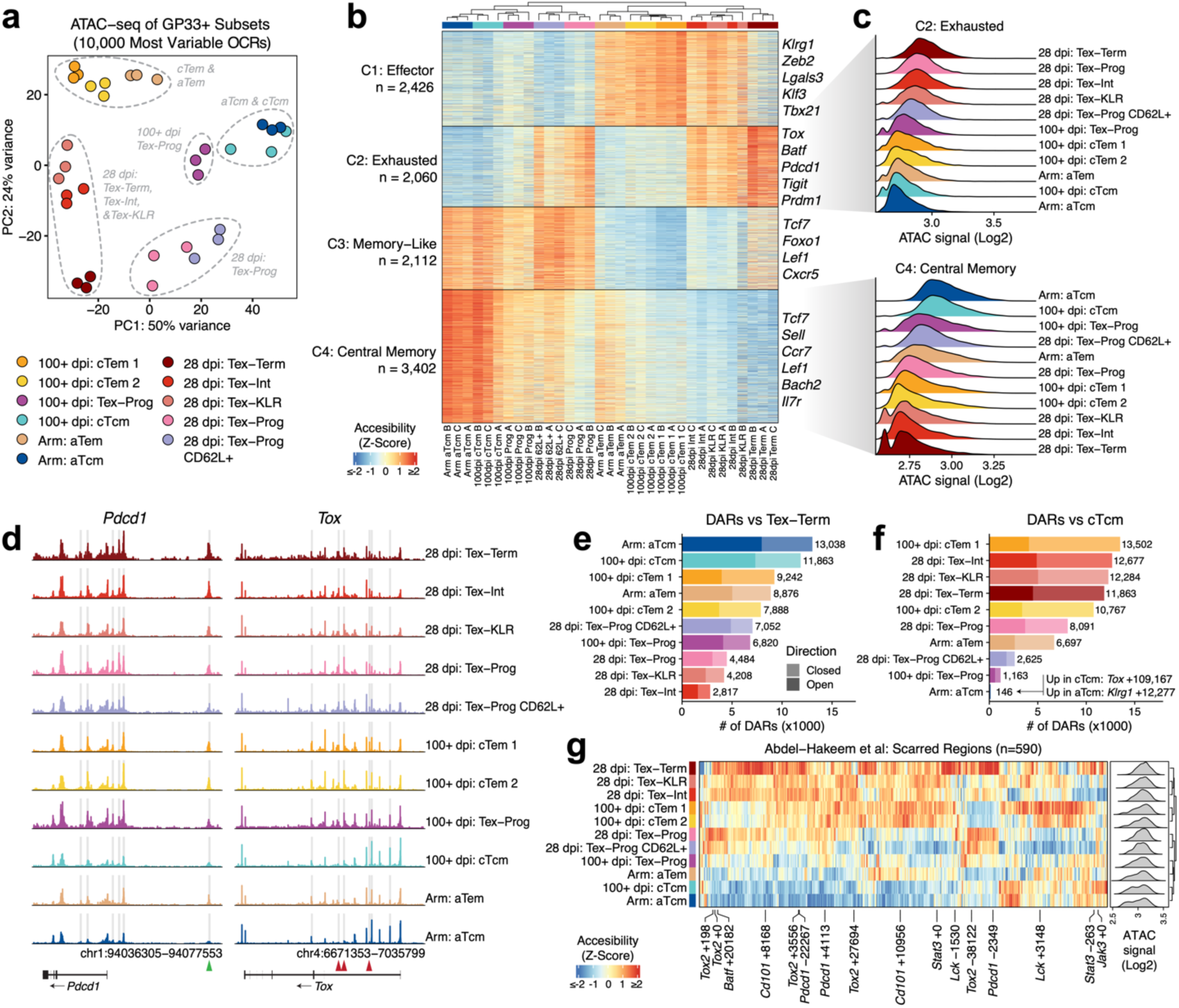
Chronic Tcm and acute Tcm have a shared epigenomic profile. ATAC-seq was performed on sorted subsets of GP33^+^ CD8^+^ T cell subsets from spleens of mice infected with LCMV-Cl13 (28 or 100+ dpi) or LCMV-Arm (100+ dpi). (a) PCA of top 10,000 most variable open chromatin regions (OCRs). Dotted lines show k-means clustering of samples based on accessibility of top 10,000 OCRs. (b) Accessibility of top 10,000 OCRs among samples. OCRs are clustered by k-means clustering. (c) Accessibility of OCRs in C2 (exhaustion-specific) or C4 (central memory-specific) across groups. (d) Genome tracks for *Pdcd1* and *Tox* across groups. Shaded boxes show the top 10,000 OCRs. Green arrow marks −23kb *Pdcd1* enhancer. Red arrows denote *Tox* OCRs in exhaustion-specific C2. (e,f) Number of differentially accessible regions (DARs) between indicated cells and Tex-Term (e) or cTcm (f). (g) Accessibility of OCRs corresponding to epigenetically scarred regions from ref^28^ across samples. (a-g) Each sample is a pool of 2-4 mice. n=3 pools per group.

We next analyzed the accessibility of OCRs across all samples to identify *cis*-regulatory elements associated with distinct cell states, and linked OCRs to genes based on the nearest transcription start site (TSS; **Fig. 3b,c**; **Supplementary Table 3**; ref^38,39^). Two clusters exhibited chromatin accessibility of effector-(C1; *Klrg1, Zeb2, Tbx21*-associated OCRs) and exhaustion-associated genes (C2; *Tox, Pdcd1, Tigit-*associated OCRs), respectively (**Fig. 3b**). The other two clusters (C3, C4) exhibited accessibility of memory-associated genes, with C4 being specifically enriched for Tcm-associated (*Sell*, *Il7r*) genes, and C3 being enriched for genes expressed by both Tcm and Tex-Prog (*Tcf7*, *Lef1*; **Fig. 3b**). Notably, compared to other subsets from cleared LCMV-Cl13, cTcm had decreased accessibility for C2 exhaustion-related OCRs, which included the exhaustion-specific −23kb *Pdcd1*-associated enhancer and three *Tox*-associated elements (*+*120kb +109kb +59kb; **Fig. 3c,d**; ref^17,40^). Similarly, cTcm had increased accessibility of C4 Tcm-related OCRs, including a *Tcf7* promoter element (−0.89kb), and *Ccr7* (TSS, +2.5kb)*, Il7r* (TSS, −44kb), and *Lef1-*associated (+2kb, −76.7kb) elements (**Fig. 3c**). To validate differences in exhaustion-associated OCRs, we calculated the number of differentially accessible regions (DARs; Log_2_FC>1, False Discovery Rate [FDR] <0.01) between Tex-Term and other subsets (**Fig. 3e**; **Supplementary Table 4**). We found that cTcm and aTcm had the most DARs when compared to Tex-Term (13,038 and 11,863, respectively), and that cTcm had decreased accessibility of seven *Tox-associated cis*-regulatory elements (e.g. TSS, *+*120kb, +109kb) and the *Pdcd1* −23kb enhancer compared to Tex-Term (**Fig. 3e**). Other populations from cleared LCMV-Cl13 showed fewer DARs versus Tex-Term than cTcm (median=7,888; **Fig. 3e**), suggesting greater epigenomic similarity to the Tex-Term cell state relative to cTcm. Conversely, by comparing cTcm to other cell subsets, we saw fewer DARs versus aTcm (n=146) than other subsets (n>1,100), again supporting the similarity between cTcm and aTcm (**Fig. 3f**). Differences in cTcm and aTcm included increased accessibility of elements associated with *Tox* (+20kb, +109kb) and *Ifi27l2a* (TSS), concordant with gene expression data from scRNA-seq, and decreased accessibility of effector-related gene elements *Klrg1* (+13kb +12kb), which was not seen at the transcriptional level (**Fig. 3f**). Together, this analysis highlights that cTcm exhibit relatively few epigenetic changes from aTcm, compared to other chronic memory cells.

To compare our results to the studies that established persistent epigenetic changes in Tex as a result of chronic stimulation (termed “epigenetic scarring”), we generated a set of overlapping OCRs with those identified by Abdel-Hakeem et al^28^. In that study, total P14 T cells were recovered from LCMV-Cl13 infected mice and allowed to rest in antigen-free hosts for at least 4 weeks (Tex-Rec). These Tex-Rec were compared with P14 from chronic LCMV-Cl13 infection (Tex) or after clearance of LCMV-Arm (Tmem) by ATAC-seq. We identified 5,781 DARs from this dataset that overlapped with OCRs in our data and compared accessibility of these regions across samples (**Extended Data Fig. 5b**). Indeed, we saw increased accessibility of OCRs associated with Tex epigenetic scarring (e.g. *Tox2* TSS, *Cd101* +8kb, *Stat3* TSS, *Batf* +20kb, *Id2* TSS) in samples from 100+ dpi of LCMV-Cl13 compared to acute memory cells, however, this signature was most pronounced in cTem and Tex-Prog compared to cTcm (**Fig. 3g**). Strikingly, cTcm displayed similar accessibility of these Tex-Rec associated OCRs as cTcm (**Fig. 3g**). Taken together, these data suggest that epigenomic changes that result from chronic stimulation may predominantly persist in cTem and persisting Tex subsets after viral clearance, rather than cTcm, which maintain a chromatin state that is highly similar to aTcm and may underlie *bona fide* memory function.

### Functional memory recall responses by cTcm

Previous studies have shown that memory-like cells that persist after chronic infection have impaired capacity to expand and differentiate upon recall response^4,27,28^. However, polyclonal memory T cell pools are comprised of individual clones with differential capacity to expand and differentiate upon rechallenge^41–43^. Based on their transcriptional and epigenetic profile, we hypothesized that cTcm biased clones would preferentially respond to rechallenge. To test clonal heterogeneity during rechallenge, we transferred polyclonal CD8^+^ T cells from mice with cleared LCMV-Cl13 or LCMV-Arm into congenic recipients and rechallenged with LCMV-Arm (**Fig. 4a**). In line with previous studies, bulk T cells transferred from cleared LCMV-Cl13 donors (Cl13:Arm) displayed decreased expansion (FC=16.3 *P*=0.0079; **Fig. 4b**) and increased PD-1^+^ frequency (FC=2.0 *P*=0.0079; **Extended Data Fig. 6b**) compared to cells from cleared LCMV-Armstrong (Arm:Arm) donors by flow cytometry^27,28^. Once we established that this model aligned with previous studies, we expanded the analysis to study the recall response of individual clones. First, we validated that characteristics of clones were replicable between hosts by evaluating expansion and phenotype among clonally-related cells challenged in five separate hosts by scRNA/TCR-seq. Cells from the same clone had significantly correlated clone size (*P*<1.1×10^-5^; **Fig. 4c**, **Extended Data Fig. 6d**) and gene expression (*P*<2.2×10^-16^; **Fig. 4c**, **Extended Data Fig. 6e**), compared to randomly generated clones, demonstrating the feasibility of studying clonal differentiation patterns of individual Cl13:Arm T cell clones during a recall response using this model.

**Figure 4:**
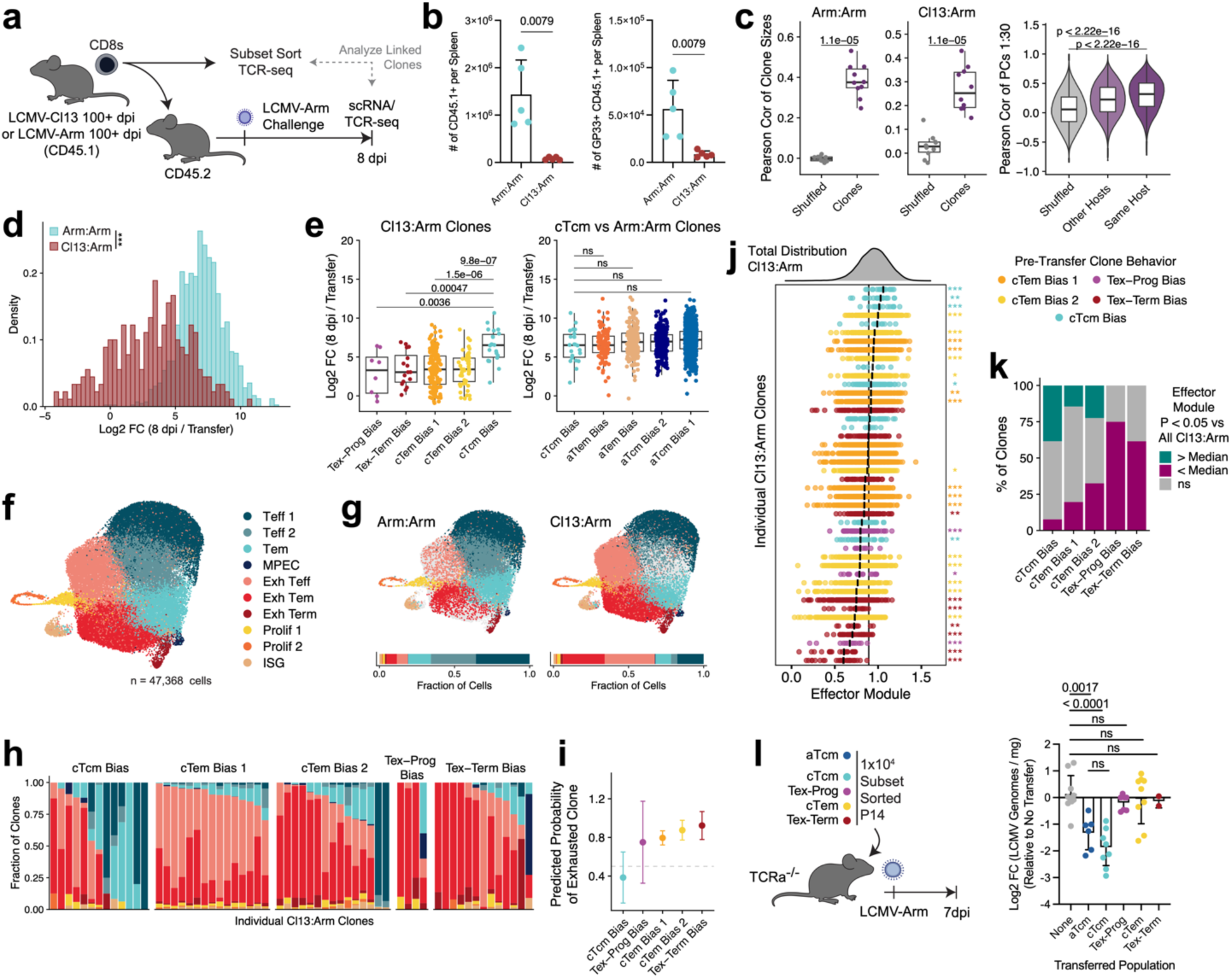
Chronic Tcm are functional upon rechallenge. (a-k) 2×10^5^ polyclonal CD45.1^+^ CD8^+^ T cells from LCMV-Arm or LCMV-Cl13 infected mice 100+ dpi were transferred into congenic hosts and challenged with LCMV-Arm. Clonal behaviors of cells pre-rechallenge determined by TCR-seq of sorted subsets (**Extended Data Fig. 6f**), and transferred cells were analyzed by scRNA/TCR-seq from the spleen 8 dpi. (b) Quantification of total and GP33^+^ transferred T cells per group. Data shown as mean+/−SD. *P* values calculated by Mann-Whitney U Test. (c) Pearson correlation of (left) clone sizes of clones between hosts and (right) gene expression PCs 1-30 of clonal sisters within the same host or in other hosts. *P* values calculated by Mann-Whitney U Test. (d-e) Clonal expansion of individual clones from (d) indicated donor or (e) clones from different pre-transfer clonal behaviors. *P* values calculated by Mann-Whitney U Test. (f, g) UMAP scRNA-seq profiles of cells 8 dpi colored by cell cluster from (f) all cells or (g) indicated group. (h) Cluster distribution of cells 8 dpi of individual Cl13:Arm clones (n>4 cells) grouped based on pre-rechallenge behavior. (i) Probability of clones from different pre-rechallenge behaviors becoming exhausted clones at 8 dpi (>50% in Exh Teff, Exh Tem, Exh Term clusters). Probability +/− 95% CI calculated by logistic regression. (j) Effector module score of cells from individual Cl13:Arm clones (n>4 cells). *P* values compare individual clones to total distribution of Cl13:Arm cells by Benjamini-Hochberg corrected Mann-Whitney U test. (k) Frequency of Cl13:Arm clones in each behavior with effector score distributions significantly above or below the total Cl13:Arm distribution. (a-k) n=5 mice/group. (l) Viral burden in the spleen of TCRα^−/−^ mice 7dpi after transfer of 1×10^4^ subset sorted P14 from LCMV-Cl13 100+ dpi or aTcm from LCMV-Arm 30+ dpi and subsequent LCMV-Arm challenge. n=10 no transfer, n=6 aTcm, n=9 cTcm, n=7 Tex-Prog, n=9 cTem, n=2 Tex-Term. Data pooled from two experiments. *P* values calculated by Mann-Whitney U Test. (a-l) Boxplots show Median, IQR, and 1.5xIQR. Barplots show mean+/−SD. *=*P*<0.05, **=*P*<0.01, ***=*P*<0.001,ns=*P*>0.05.

We next evaluated the clonal expansion of individual clones after transfer and observed an overall decrease in clonal expansion among Cl13:Arm (Median=3.36) cells compared to Arm:Arm cells (Median=7.04; *P*<2.2×10^-16^; **Fig. 4d**). Surprisingly, a subset of Cl13:Arm clones (20/261 clones) expanded to the same degree as Arm:Arm clones (≥Arm:Arm median; **Fig. 4d**), suggesting that while the total pool of antigen-specific cells after chronic antigen clearance has impaired recall capacity, there are individual clones that retain functional memory capacity. To determine the origin of these functional chronic antigen-specific clones, we identified clonal behaviors pre-transfer by sorting T cell subsets and performing bulk TCR-seq (**Extended Data Fig. 6f**). Consistent with our prior analysis, we identified five clonal behaviors of Cl13:Arm cells pre-rechallenge by clustering: cTcm bias, Tex-Prog bias, Tex-Term bias, and two cTem bias clusters (**Extended Data Fig. 6f**). We assessed the expansion of clones in each behavior after rechallenge and observed that cTcm biased clones expanded significantly better than Cl13:Arm clones from other pre-transfer behaviors (cTem Bias 1: FC=6.31 *P=*1.5×10^-6^, cTem Bias 2: FC=7.75 *P=*9.8×10^-7^, Tex-Prog Bias: FC=8.98 *P=*0.0036, Tex-Term Bias: FC=7.17 *P=*0.00047; **Fig. 4e**). Moreover, there was no significant difference in expansion between cTcm biased clones and Arm:Arm clones, including aTcm biased clones, (*P*>0.05; **Fig 4e**; clonal behaviors of Arm:Arm clones determined by the same method as Cl13:Arm; **Extended Data Fig. 6f**), demonstrating the ability of cTcm-based clones to robustly expand upon rechallenge to the same degree as acute memory cells.

To further support these findings, we evaluated the recall responses of cTcm *in vivo* using cell transfer experiments. cTcm and other subsets of P14 cells from 100+ dpi of LCMV-Cl13 (cTem, Tex-Prog, and Tex-Term) or P14 aTcm from 50+ dpi of LCMV-Arm were transferred into naive congenic hosts and challenged with LCMV-Arm (**Extended Data Fig. 7a,b**). At 7 dpi, we quantified the expansion of P14 cells by flow cytometry and found that there were significantly higher numbers of cTcm-derived cells compared to other chronic memory subsets (Tex-Prog: FC=1.77 *P*=0.0104, cTem: FC=50.6 *P*<1×10^4^, Tex-Term: FC=105.1 *P*<1×10^4^; **Extended Data Fig. 7c**). Importantly, cTcm expanded and differentiated into KLRG1^+^ effectors to a similar degree as aTcm (*P*>0.05 ; **Extended Data Fig. 7c,d**), although a significantly increased number of cTcm were PD-1^+^ upon rechallenge compared to aTcm (FC=3.75 *P*=0.025; **Extended Data Fig. 7d**). To assess the capacity of individual Cl13:Arm clones to differentiate into effector cells after rechallenge, we analyzed scRNA/TCR-seq (n=47,369 cells total) of transferred CD8^+^ T cells and endogenous GP33^+^ cells from 8 dpi with LCMV-Arm (**Fig. 4f**). scRNA/TCR-seq clustering revealed 10 clusters of cells, which were identified by DEGs (LogFC>0.25; **Fig. 4f**; **Supplementary Table 5**). These included effector cells (Teff; expressing *Zeb2, Klrg1*), memory-precursor effector cells (MPEC; expressing *Tcf7, Ltb*), and Tem (expressing *Cxcr3, Il7r*) clusters, as well as exhaustion-associated clusters, characterized by *Tox* and *Pdcd1* expression along with Teff (*Gzma, Cx3cr1*), Tem (*Cxcr3, Il7r*), or Tex-Term (*Tigit, Lag3, Cd7*) genes (**Extended Data Fig. 7e**). In line with flow cytometry data, the majority of Arm:Arm and endogenous GP33^+^ cells were in Tem and Teff clusters, while most Cl13:Arm cells were in exhausted clusters (**Fig. 4g**, **Extended Data Fig. 7f**). However, 32.3% of Cl13:Arm cells did not appear phenotypically exhausted and were in Teff and Tem clusters (**Fig. 4g**). At the clonal level, we found a subset of clones (n=39/187) that had a low proportion of cells in the exhaustion-associated clusters (**Fig. 4h**). We assessed the origins of these clones and determined that the probability of cTcm biased clones becoming >50% exhausted (38.5% +/−26.4% CI by logistic regression) was lower than clones from other Cl13:Arm behaviors (>75% probability; **Fig. 4i**). Similarly, cTcm biased clones had lower expression of genes including *Tox*, *Pdcd1*, and *Lag3*, relative to other Cl13:Arm clones (**Extended Data Fig**). Next, we tested the ability of individual clones to upregulate effector related genes (including *Klrg1, Ccr2, Ly6c2*; **Supplementary Table 7**) and found that some clones had a significantly higher effector module score than the total population of Cl13:Arm cells (**Fig. 4j**). Overall, 38.5% of cTcm biased clones had a significantly higher effector module score than the total population of Cl13:Arm cells, a higher frequency than any other clone behavior (**Fig. 4j,k**). As validation, we observed increased expression of individual genes (*Klrg1, Ccr2, Ly6c2*) in the effector gene module among cTcm clones compared to other Cl13:Arm clones (**Extended Data Fig 7g**). Taken together, these data suggest that cTcm biased clones can robustly expand and differentiate into effector cells upon rechallenge.

Finally, to assess whether increased clonal expansion and differentiation enables pathogen clearance by cTcm, we transferred P14 T cell subsets into TCR⍺^−/−^ mice, which lack endogenous CD4^+^ and CD8^+^ T cells, and challenged with LCMV-Arm. Transfer of either aTcm or cTcm led to significant reduction in LCMV viral burden compared to untreated controls (aTcm: FC=2.22 *P=*0.0017; cTcm: FC=3.35 *P*<0.0001), while other subsets of CD8^+^ chronic memory cells had no impact on viral burden (*P*>0.05; **Fig. 4l**). Importantly, there was no significant difference between cTcm and aTcm (*P*>0.05), further demonstrating the functional recall response of cTcm (**Fig. 4l**). Together, these findings demonstrate that the expansion and differentiation of cTcm biased clones upon rechallenge enables protective immunity.

### ⍺PD-L1 treatment expands cTcm biased clones

Since Tex-Prog preferentially respond to ICB and cTcm are derived from Tex-Prog biased clones, we assessed the impact of ICB on cTcm. Specifically, we tested whether ICB: (1) impacted the durability of cTcm, and (2) led to the expansion of cTcm biased clones^10,15,16^. To test this, we treated LCMV-Cl13 infected mice with a monoclonal antibody blocking PD-L1 (⍺PD-L1), the ligand for PD-1 (**Fig. 5a**). As shown previously, ⍺PD-L1 treatment led to significant expansion of GP33^+^ cells during the chronic phase of infection (FC=1.62 *P*=0.028; **Fig. 5b**); however, there were no significant (*P*>0.05) differences in GP33^+^ frequency or count after viral clearance between groups (**Fig. 5b**, **Extended Data Fig. 8a;** ref^44^). We performed scRNA/TCR-seq on splenic GP33^+^ T cells at 100+ dpi to assess if ⍺PD-L1 impacted the frequency or gene expression profiles of chronic memory cells and again found no significant changes in subset frequencies of chronic memory T cells after ⍺PD-L1 treatment, compared to PBS-treated controls (**Fig. 5c**, **Extended Data Fig. 8b-d**, **Supplementary Table 6**), which we further validated with flow cytometry (**Extended Data Fig. 8e**). Within cell clusters, gene expression was also highly correlated (R>0.9, *P*<0.001) between groups and there were less than 10 DEGs between groups across all clusters (**Extended Data Fig. 8f,g**). Finally, we assessed if clonal behaviors were altered by ⍺PD-L1 treatment, and found all clonal behaviors, including cTcm biased clones, in ⍺PD-L1 treated mice (**Fig 5d**).

**Figure 5:**
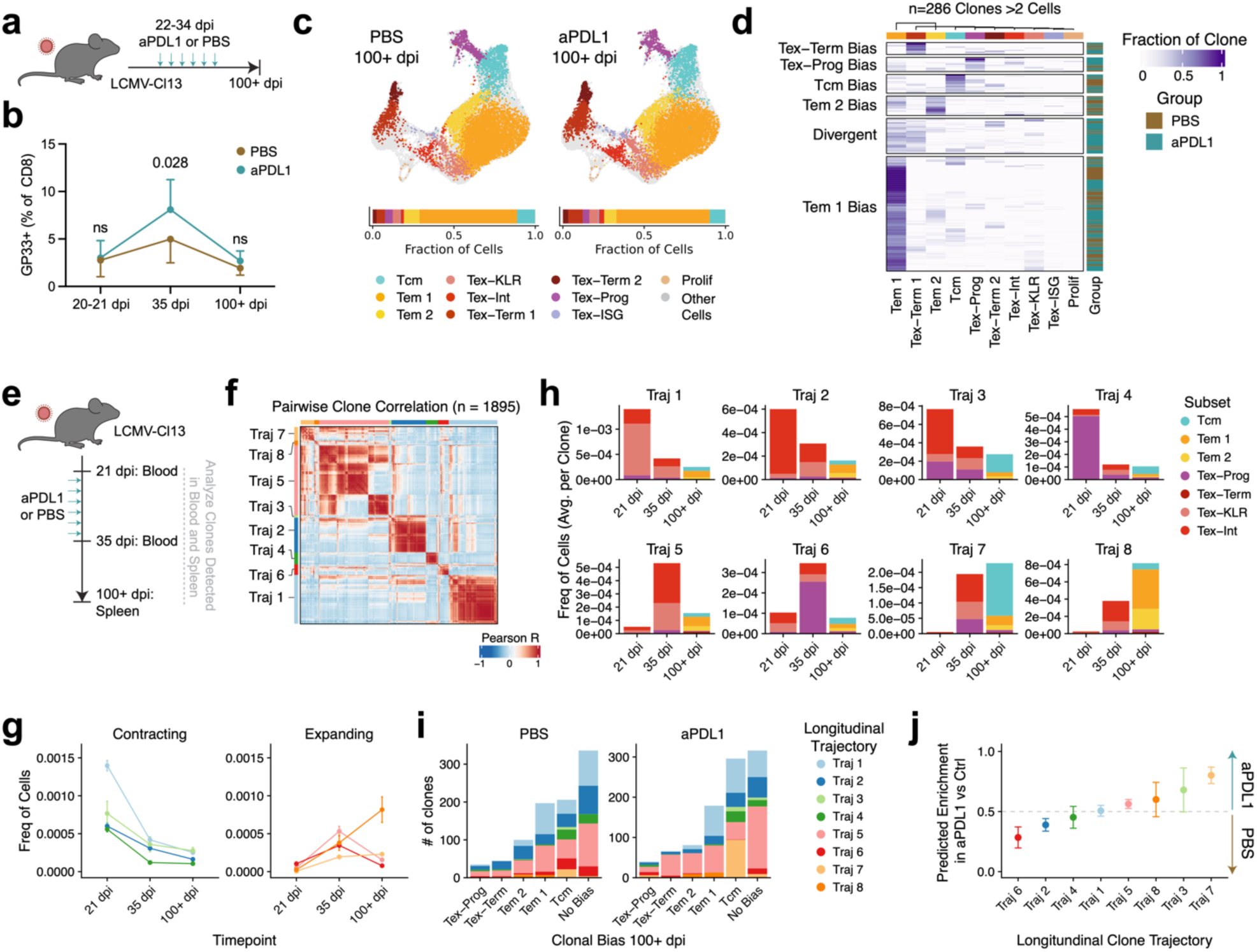
*⍺*PD-L1 treatment expands chronic Tcm biased clones. (a) Mice were infected with LCMV-Cl13 and treated with ⍺PD-L1 from 22 to 34 dpi. (b) Flow cytometry of GP33^+^ cell frequency from blood (pre-treatment; 20-21 dpi and post-treatment; 35 dpi) or spleen (100+ dpi). Mean +/− SD. n=12 mice per group from two independent experiments. *P* values calculated by Mann-Whitney U Test. (c-d) 100+ dpi, scRNA/TCR-seq was performed on GP33^+^ cells from the spleen and integrated with data from Fig. 1. (c) Cells from PBS or ⍺PD-L1 treated mice colored by cell cluster. Cells in grey are from other datasets. (d) Phenotype distribution of individual TCR clones across subsets, grouped by clonal behavior determined by k-means clustering. (c-d) n=4 PBS mice, n=5 ⍺PD-L1 mice. (e-j) Cell subsets were sorted from the blood (21 and 35 dpi) or spleen (100+ dpi) of LCMV-Cl13 infected mice treated with PBS or ⍺PD-L1. Bulk TCR-seq was used to trace clonal differentiation across timepoints. (f) Pairwise correlations of clone subset frequencies at each timepoint. Common longitudinal trajectories (Traj) were identified by clustering based on correlations. (g) Clone size of clones in each longitudinal trajectory over time. Plot shows mean +/− SEM. (h) Average subset frequency (of all CD8^+^ T cells) for clones in each trajectory at different timepoints. (i) Number of clones per group that result in clones biased towards each phenotype at 100+ dpi. Bias determined by *P*<0.05 by hypergeometric test with Hochberg correction. (j) Enrichment of clonal trajectories in ⍺PD-L1 group vs PBS group. Probability +/− 95% CI calculated by logistic regression. (e-j) n=8 PBS mice, n=7 ⍺PD-L1 mice.

Given that cTcm preferentially differentiate from Tex-Prog biased clones, we hypothesized that ⍺PD-L1 treatment may selectively expand clones that form cTcm. Moreover, ICB acts by expanding pre-existing Tex-Prog clones as well as by priming new clones, so we tested the impact of both mechanisms of action on the formation of cTcm^45,46^. To evaluate the contribution of ⍺PD-L1 mobilized clones to the cTcm pool, we performed longitudinal bulk TCR-seq on T cell subsets from blood pre- and post-⍺PD-L1 treatment (Tex-Prog, Tex-Int, and Tex-KLR), and at a memory timepoint (Tcm, Tem 1, Tem 2, Tex-Prog, and Tex-Term; **Fig. 5e**, **Extended Data Fig. 4c**). Again, we did not restrict analysis to GP33^+^ clones, because T cells with low avidity TCRs preferentially respond to ICB^47^. To assess clonal trajectories over time, we adapted an existing analytical framework (Cyclone; ref^48^) to determine longitudinal trajectories based on the clustering of pairwise correlations of clone subset frequencies at each timepoint (**Fig. 5f**). This approach yielded eight clonal trajectories, including clones that contracted over time (Trajectory 1-4) and clones that expanded after the treatment period (Trajectory 5-8; **Fig. 5f,g**). Expanding clonal trajectories included clones not present before treatment, which could either be *de novo* primed clones or clones present at low/undetectable frequencies prior to ⍺PD-L1. Utilizing longitudinal phenotype information, we identified clones that phenotypically transitioned from Tex-Int and Tex-KLR to cTem (Trajectories 1,2,5,8) as well as clones that transition from Tex-Prog that resulted in cTcm (Trajectories 3,4,6,7; **Fig 5h**). We assessed which longitudinal trajectories could lead to clones with a significant bias toward cTcm (*P*<0.05 hypergeometric test), and found that cTcm biased clones could arise from all longitudinal trajectories (**Fig 5i**), as shown in earlier clone tracing experiments. Importantly, ⍺PD-L1 treatment yielded more cTcm biased clones, primarily from clones in Trajectory 7 (FC=2.97) and few cTcm biased clones from Trajectory 6 (FC=41.7; **Fig. 5j**). Trajectory 7 was enriched for clones that were small or undetected pre-treatment, evenly spread across phenotypes at d35, further expanded from 35 dpi to 100 dpi, and primarily formed majority cTcm (**Fig. 5g,h**). Trajectory 6 clones were generally >70% Tex-Prog at 35 dpi, formed cTcm at 100+ dpi, and contracted from 35 to 100+ dpi (**Fig. 5g,h**). Among all clone trajectories, Trajectory 7 was the most enriched in the ⍺PD-L1 group compared to control (80.2% +/−3.48% CI in ⍺PD-L1 by logistic regression), while Trajectory 6 was depleted (28.4% +/−4.47% CI in ⍺PD-L1; **Fig. 5j**). These data suggest that ⍺PD-L1 ICB may drive the proliferation and differentiation of previously unexpanded T cell clones that contribute to the cTcm pool after antigen clearance.

## Discussion

Here, we describe a clonally restricted subset of Tex that maintains the phenotypic plasticity to form functional long-lived memory T cells. We utilized clonally resolved phenotypic profiling of the polyclonal T cell response to uncover multiple differentiation trajectories of T cells that persist following chronic antigen clearance. Specifically, cTcm preferentially differentiated from Tex-Prog biased clones, and cTem developed from more differentiated effector-like Tex-Int and Tex-KLR clones, mirroring analogous memory differentiation trajectories after acute infection. cTcm exhibited transcriptional and epigenetic states that were nearly identical to aTcm, and they were able to robustly expand, differentiate into effector T cells, and clear secondary viral infections to the same extent as aTcm.

Prior studies have shown that T cells maintain exhaustion-related epigenetic changes or “scars” from chronic stimulation that impair their subsequent recall response^25,27,28^. Our data are consistent with this finding as we observe that the majority of Tex-derived cells–including cTem and Tex subsets–maintain chromatin accessibility of exhaustion-associated loci after clearance of chronic viral infection, and accordingly, are unable to respond to rechallenge. However, we find that a subset of Tex-Prog clones is able to avoid this epigenetic fate and lead to cTcm. Since cTcm are a relatively small fraction of the total antigen-specific T cell pool after viral clearance, we suspect that the functional response and genomic signatures of these clones may be masked in population-level analyses. Previous work has suggested that the preservation of Tex-Prog in the setting of chronic antigen can be maintained by cell extrinsic factors, including the sequestration of T cells in low antigen density spatial niches^9,36,49,50^. Our results may support and extend this concept, since chronic LCMV infection can persist long-term locally in many organs even after peripheral clearance; therefore, the generation of Tex-Prog biased clones and cTcm may require stable lymphoid niches that preserve memory plasticity by avoiding antigen exposure and TCR stimulation^1^. In line with this concept, low TCR signal strength–via low avidity or decreased dendritic cell activation–has been shown to promote a stem-like Tex cell state, while high avidity TCRs preferentially form Tex in both tumors and chronic infection^11,47,50–53^. This model is further supported by the robust expansion of cTcm clones after ⍺PD-L1 treatment, since low avidity clones are preferentially engaged by ICB^47^.

Our findings also complement recent studies that demonstrate parallel differentiation trajectories of stem-like T cells at the earliest stages of chronic and acute infection^54,55^. These studies have shown that: (1) T cells with high TCR avidity for antigen preferentially differentiate into Tex-Prog during acute stimulation, and (2) stem-like T cells early in infection are nearly identical in both viral challenge settings, which may represent an anticipatory mechanism to prepare the host for different infection outcomes^54,55^. Our work extends this model and demonstrates that a similar parallel exists at memory time points after pathogen clearance, where some Tex-Prog-derived cells maintain durable epigenetic adaptations to chronic stimulation, while a clonally-distinct subset of Tex-Prog-derived cells form cTcm that can robustly respond to acute secondary rechallenge, which may represent a similar anticipatory evolutionary adaptation for a broad range of secondary infection outcomes.

Finally, the expansion of cTcm biased clones upon ⍺PD-L1 treatment may provide a mechanistic explanation for several clinical observations. First, tumor-specific T cell clones that expand after ⍺PD-(L)1 treatment in patients can be found in the blood and persist for years after therapy^24,56^. However, since these clones were exhausted in the tumor microenvironment, it has been unclear if they could contribute to the memory response and promote long-term survival, which can last for over a decade^57^. Our prior work demonstrated that the presence of tumor-specific Tex-Prog in tumor-draining lymph nodes of these patients was associated with a greater likelihood of long-term persistence in the blood, and our current study bridges this gap and suggests that a fraction of the Tex-Prog population may lead to the differentiation of *bona fide* memory cells long-term^24^. Second, ⍺PD-(L)1 ICB enhances vaccine induced antitumor responses, which may augment long-term survival^58,59^. Our data suggest that this may be driven by the invigoration of novel and lowly expanded T cell clones, which contribute to the functional memory T cell repertoire and support long term-immunity. Finally, our results support growing evidence that clonal responses to ICB can be derived from two sources: (1) clonal reinvigoration or revival, where ICB leads to differentiation of pre-existing clones, and (2) clonal replacement, where ICB leads to *de novo* priming of naive T cell clones^45,46^. We find that both pre-existing and novel clones comprise the cTcm pool. Pre-existing ICB-activated clones produce differentiated Tex immediately following treatment, while also maintaining cells in their capacity to form memory. In addition, ICB leads to the expansion of novel clones that form cTcm, which may be enriched for low avidity clones that are primed due to increased TCR signaling with ICB. Altogether, these findings underscore the capacity of select exhausted T cell clones to maintain phenotypic plasticity and transition to functional memory cells, providing a foundation for future interventions to enhance immune memory to chronic infections and cancer.

## Extended Data Figures

**Extended Data Figure1:**
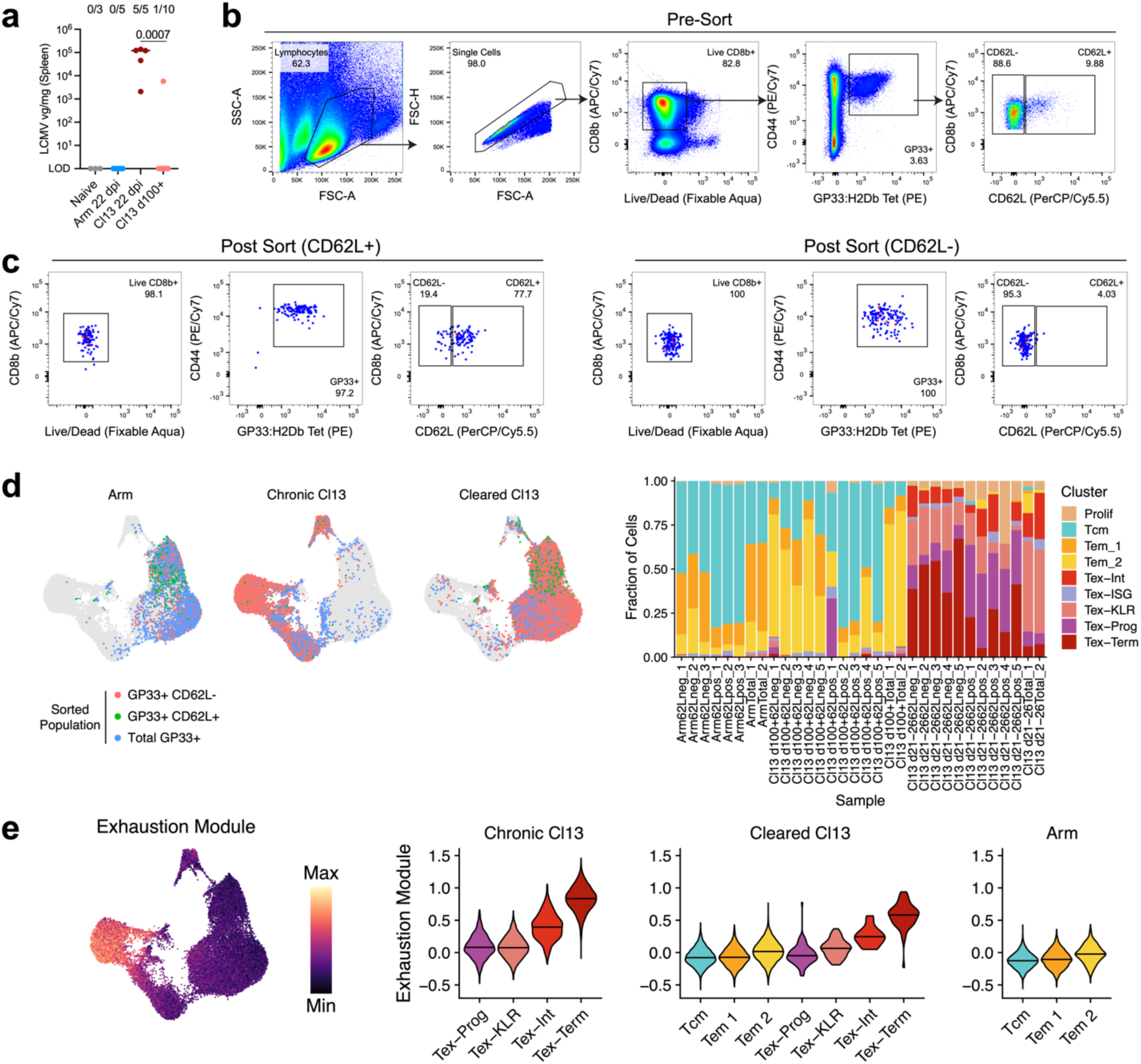
scRNA/TCR-seq categorization of GP33^+^ cells after peripheral LCMV-Cl13 clearance. (a) Viral burden in the spleen of naïve mice or mice infected with LCMV-Cl13 (22 or 100+ dpi) or LCMV-Arm (22 dpi). Lines show median. Numbers show the fraction of mice with detectable viral load. *P* value calculated by Mann-Whitney U Test. n=3 naïve, n=5 LCMV-Arm, n=5 LCMV-Cl13 22 dpi, n=10 LCMV-Cl13 100+ dpi. (b,c) Sorting scheme (b) and representative post-sort purity (c) for scRNA/TCR-seq. (d) (left) scRNA-seq UMAP of cells sorted from each timepoint colored by sort. (right) Cell cluster distribution from individual samples. (e) Exhaustion module score of all cells (left) and cells from select clusters in each group. Horizontal lines show the median.

**Extended Data Figure2:**
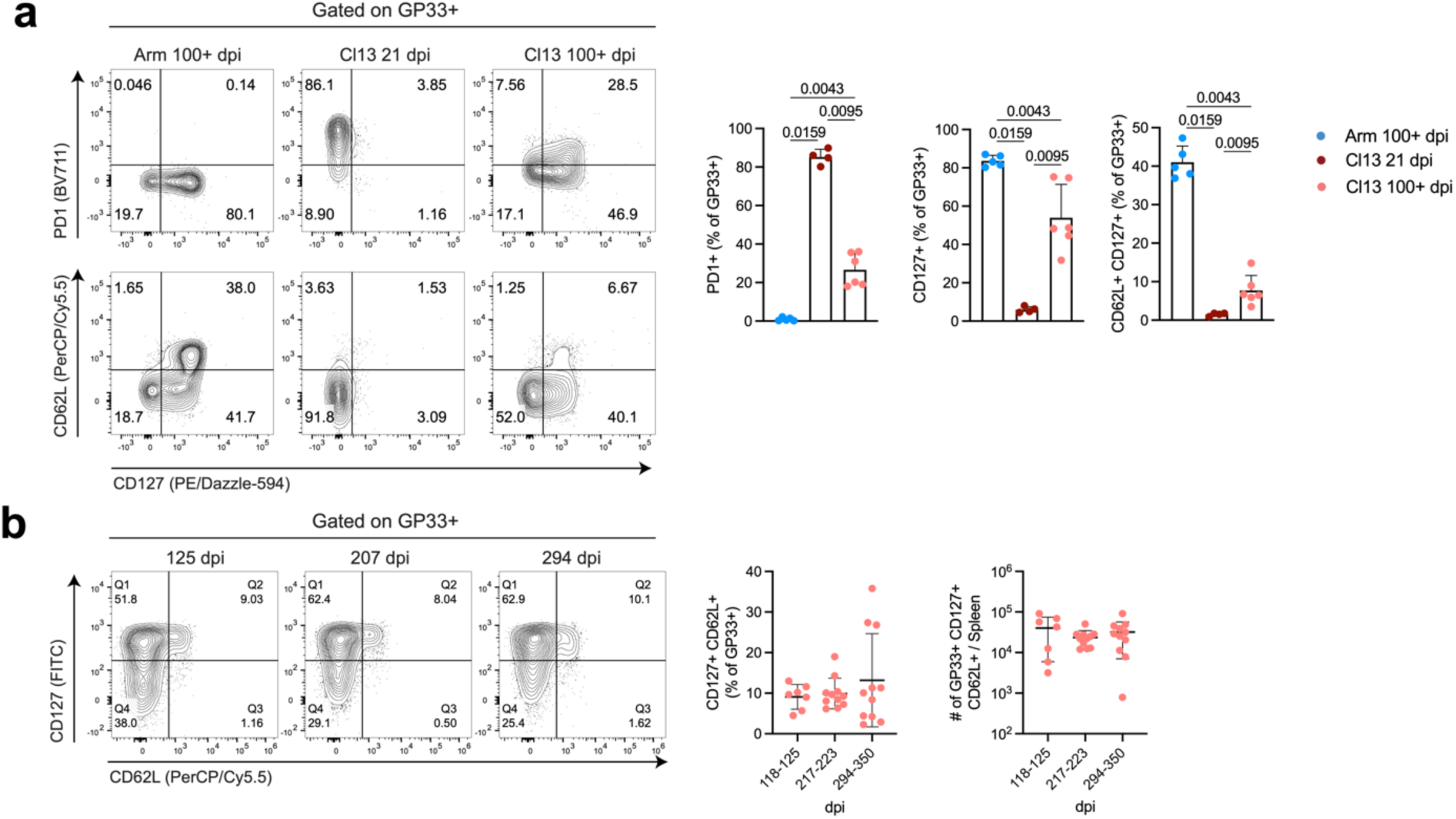
Flow cytometry of CD62L^+^ CD127^+^ Tcm after peripheral clearance of LCMV-Cl13. (a) Flow cytometry of PD-1, CD127, and CD62L among GP33^+^ CD8^+^ T cells in the spleen. n=6 LCMV-Cl13 −100+ dpi, n=5 LCMV-Arm, n=4 LCMV-Cl13 21dpi. Representative experiment of multiple independent experiments. *P* values calculated by Mann-Whitney U Test. (b) Quantification of CD127^+^ CD26L^+^ GP33^+^ cells at indicated timepoints after LCMV-Cl13 infection. n=7 118-125 dpi, n=11 217-223 dpi, n=11 294-350 dpi. Data pooled from two independent experiments. (a,b) Data shown as mean+/− SD.

**Extended Data Figure3:**
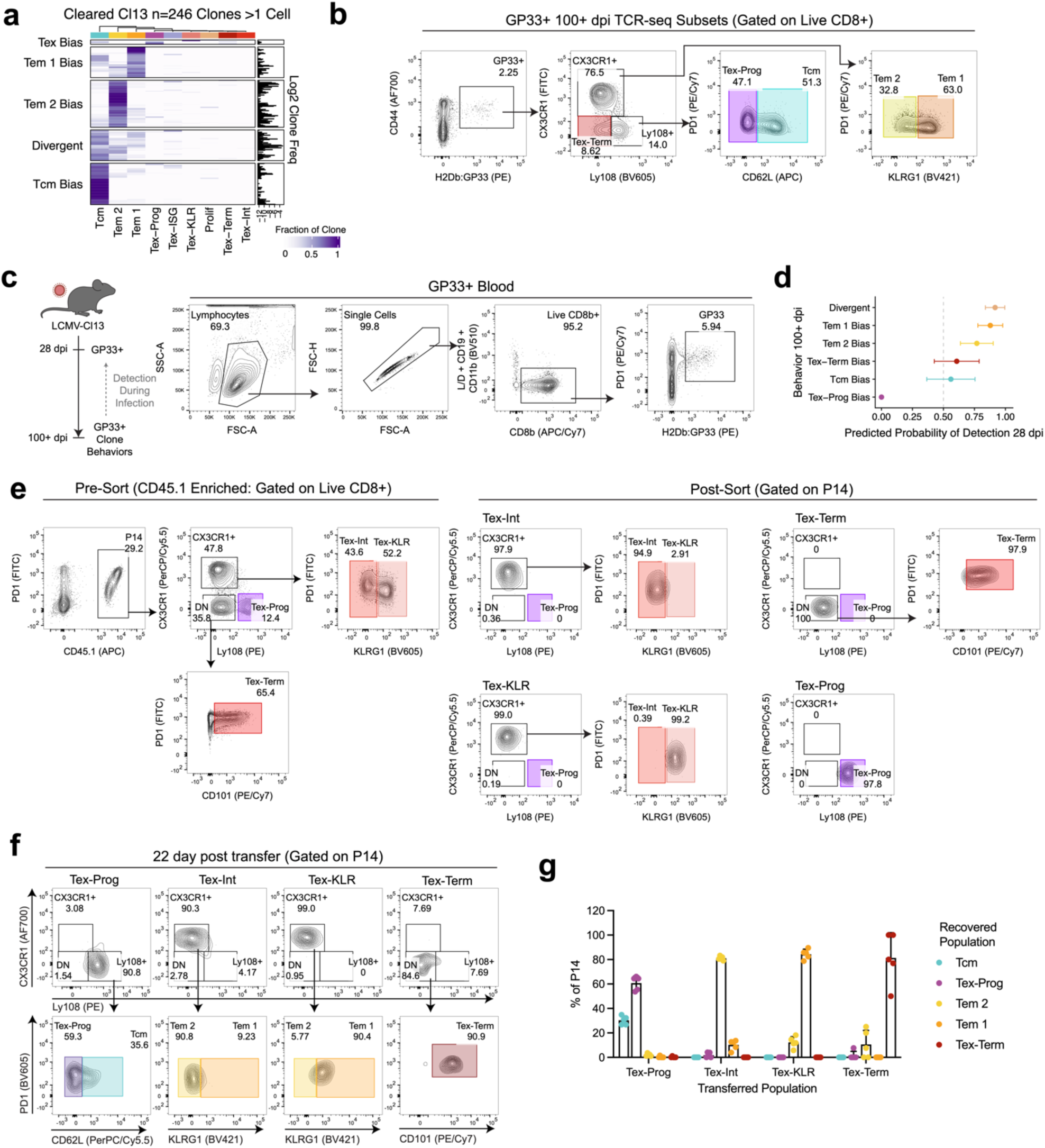
cTcm are derived from Tex-Prog. (a) Custer distribution of individual expanded clones from scRNA/TCR-seq from 100+ dpi LCMV-Cl13. Clones are grouped by clonal behavior determined by k-means clustering. (b) Gating scheme for bulk TCR-seq at 100+ dpi. (c,d) Detection of GP33^+^ clones in blood 28dpi. Clonal behaviors of GP33^+^ were determined by TCR-seq 100+ dpi (Fig. 2b) and detection during infection was tested by TCR-seq of blood GP33^+^ cells 28 dpi. (d) Probability of clones from different behaviors being detected in blood at 28 dpi. Probability +/− 95% CI calculated by logistic regression. (e-g) 4×10^3^ naive CD45.1^+^ P14 T cells were transferred into congenic primary hosts infected with LCMV-Cl13. Subsets of exhausted P14 cells were isolated and 4.5×10^4^ cells were transferred into infection free secondary hosts. After 22 days, cells were isolated and assessed by flow cytometry. (e) Sorting scheme (left) for exhausted P14 subsets and (right) sort purity. (f,g) Representative flow cytometry (f) and (g) quantification of cells recovered in secondary hosts 22 days post transfer. Data shown as mean+/−SD. n=5 mice per group. Representative experiment of two independent experiments. *P* values calculated by Mann-Whitney U Test.

**Extended Data Figure4:**
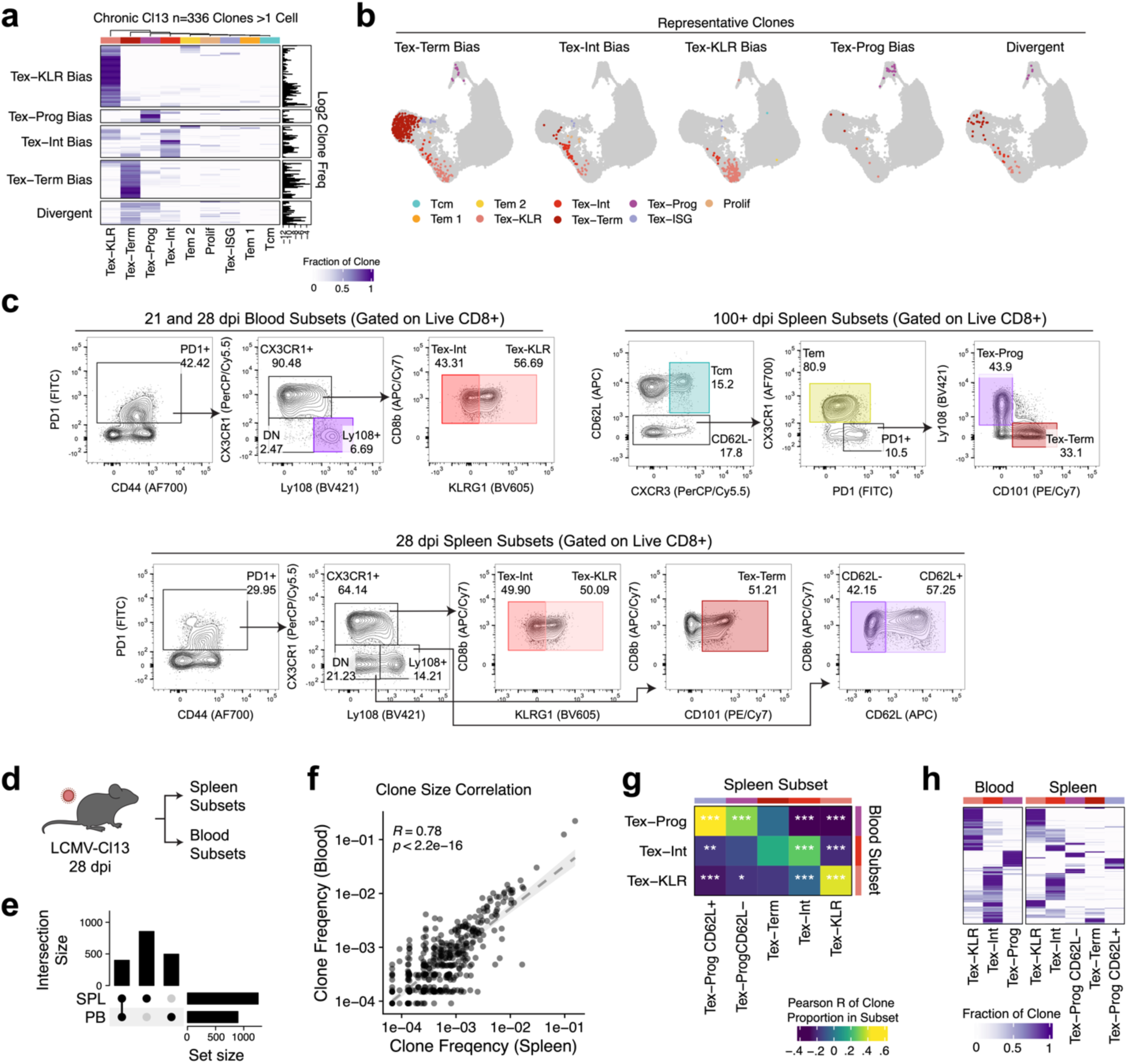
Presence of Tex-Prog biased clones in chronic infection and model validation for longitudinal clone tracing. (a,b) scRNA/TCR-seq of T cell clones from 21-26 dpi LCMV-Cl13. (a) Custer distribution of individual expanded clones from scRNA/TCR-seq. Clones are grouped by clonal behavior determined by k-means clustering. (b) Representative clones from each behavior. (c) Gating schemes for longitudinal bulk TCR-seq studies. (d-h) PD-1^+^ exhausted cell subsets were sorted from the blood of an LCMV-Cl13 infected mouse 28 dpi. (e) Upset plot of clones detected in the spleen, blood, or both organs. (f) Pearson correlation of total clone frequency in the blood and spleen. (g) Pearson correlation of clone frequency in each subset between organs. Significant correlations by Pearson correlation are highlighted (Bonferonni corrected *P* values: *=*P*<0.05, **=*P*<0.01, ***=*P*<0.001). (h) Phenotype distribution of clone frequency of shared clones in blood or spleen.

**Extended Data Figure5:**
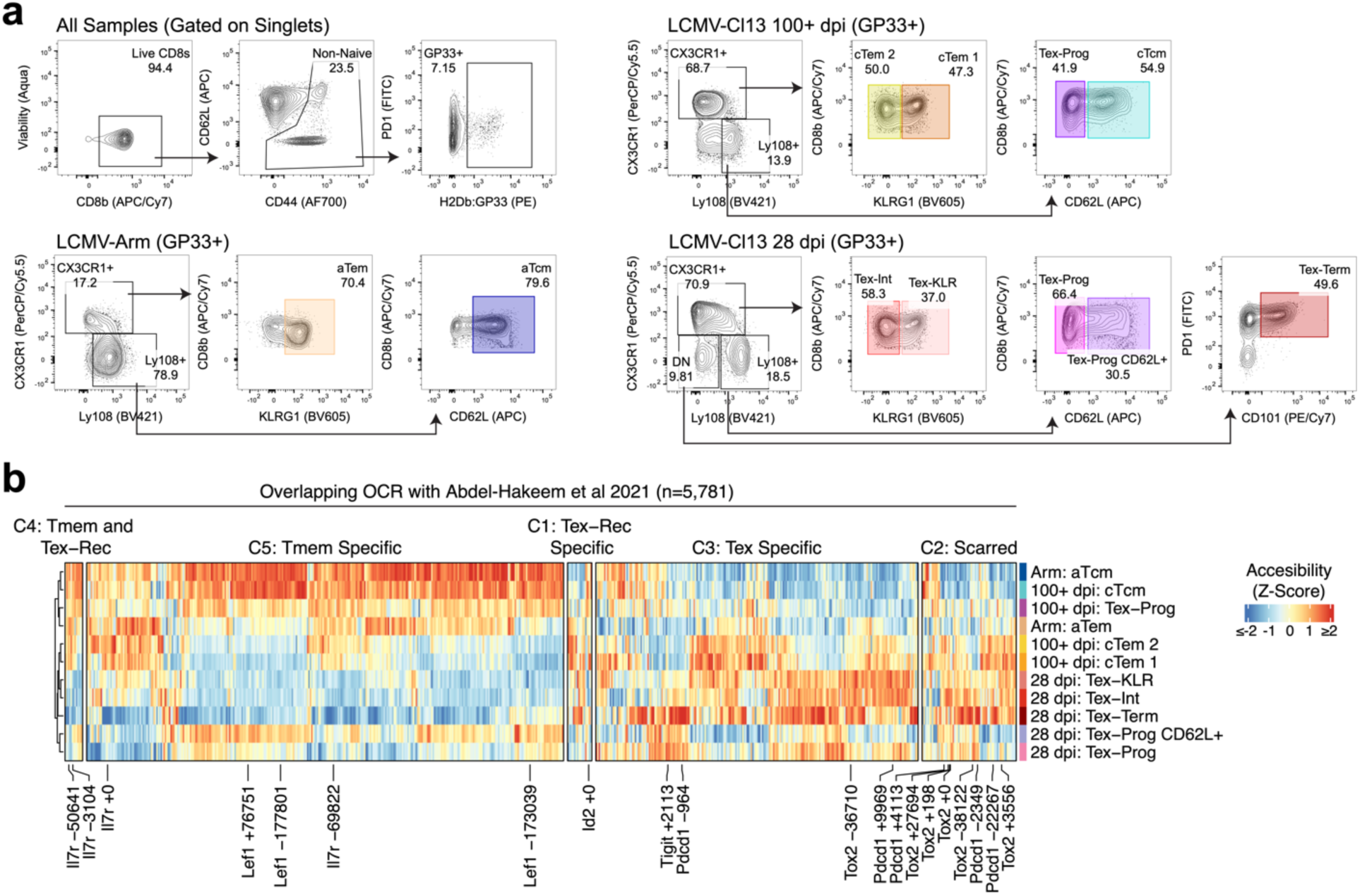
ATAC-seq gating schemes and analysis of overlapping OCRs with Abdel-Hakeem et al. (a) Gating schemes for ATAC-seq. (b) Accessibility of OCRs corresponding to regions identified in ref^28^ across samples.

**Extended Data Figure6:**
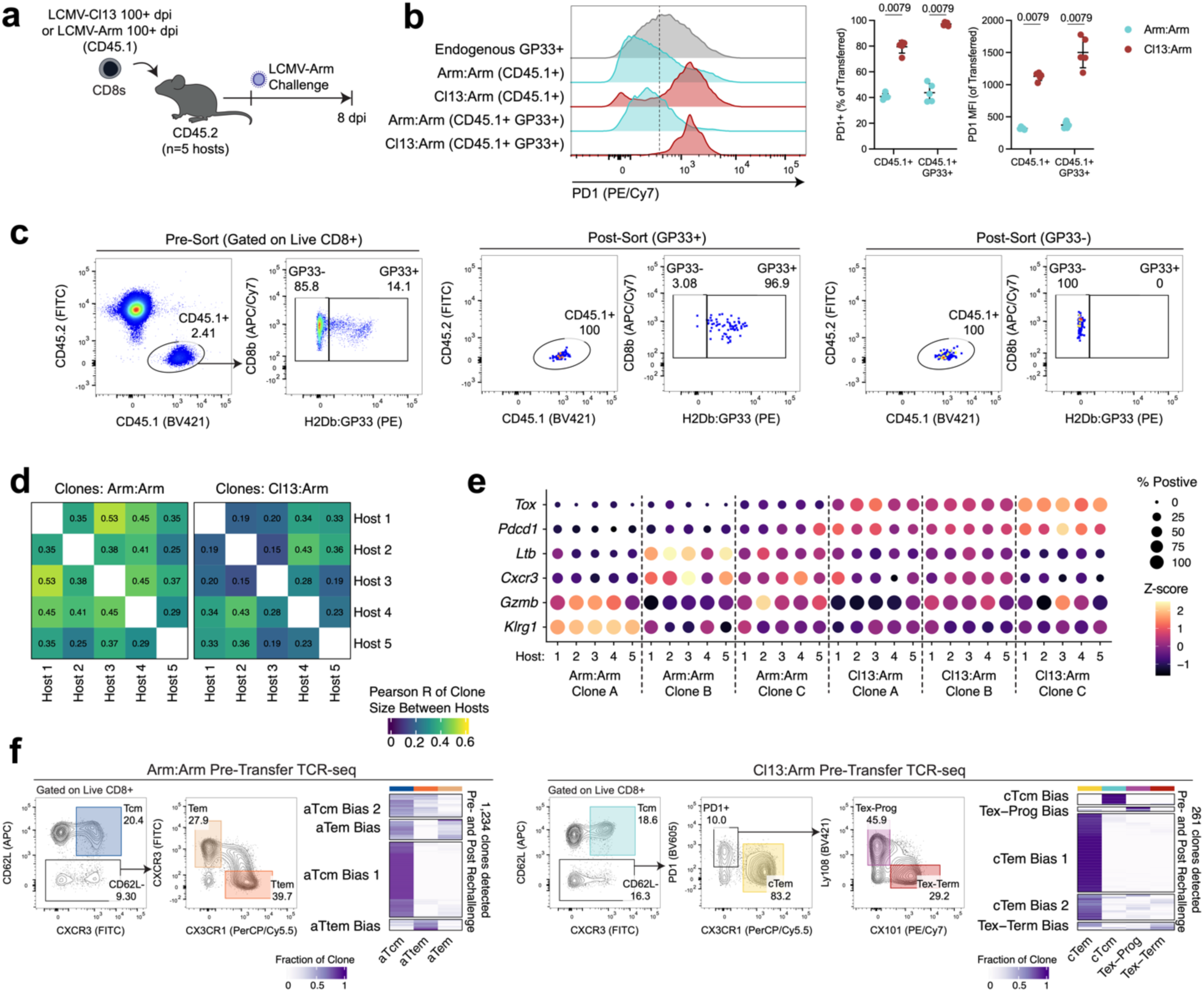
Polyclonal rechallenge of CD8^+^ T cells from 100+ dpi LCMV-Arm or LCMV-Cl13. (a-f) 2×10^5^ polyclonal CD45.1^+^ CD8^+^ T cells from LCMV-Arm or LCMV-Cl13 infected mice 100+ dpi were transferred into congenic hosts and challenged with LCMV-Arm. (b) PD-1 expression among transferred CD45.1^+^, CD45.1^+^ GP33^+^, or endogenous CD45.2^+^ GP33^+^ T cells at 8 dpi. (c) Sorting scheme and purity for scRNA/TCR-seq 8 dpi. (d) Pearson of clone sizes between individual hosts. (e) Expression of Tex (*Tox, Pdcd1*), Tem (*Ltb, Cxcr3*), and Teff (*Gzmb, Klrg1*) genes among clonally-related cells split by host. (f) Pre-rechallenge bulk TCR-seq of subset sorted T cells. Heatmaps are split by clone behavior determined by k-means clustering.

**Extended Data Figure7:**
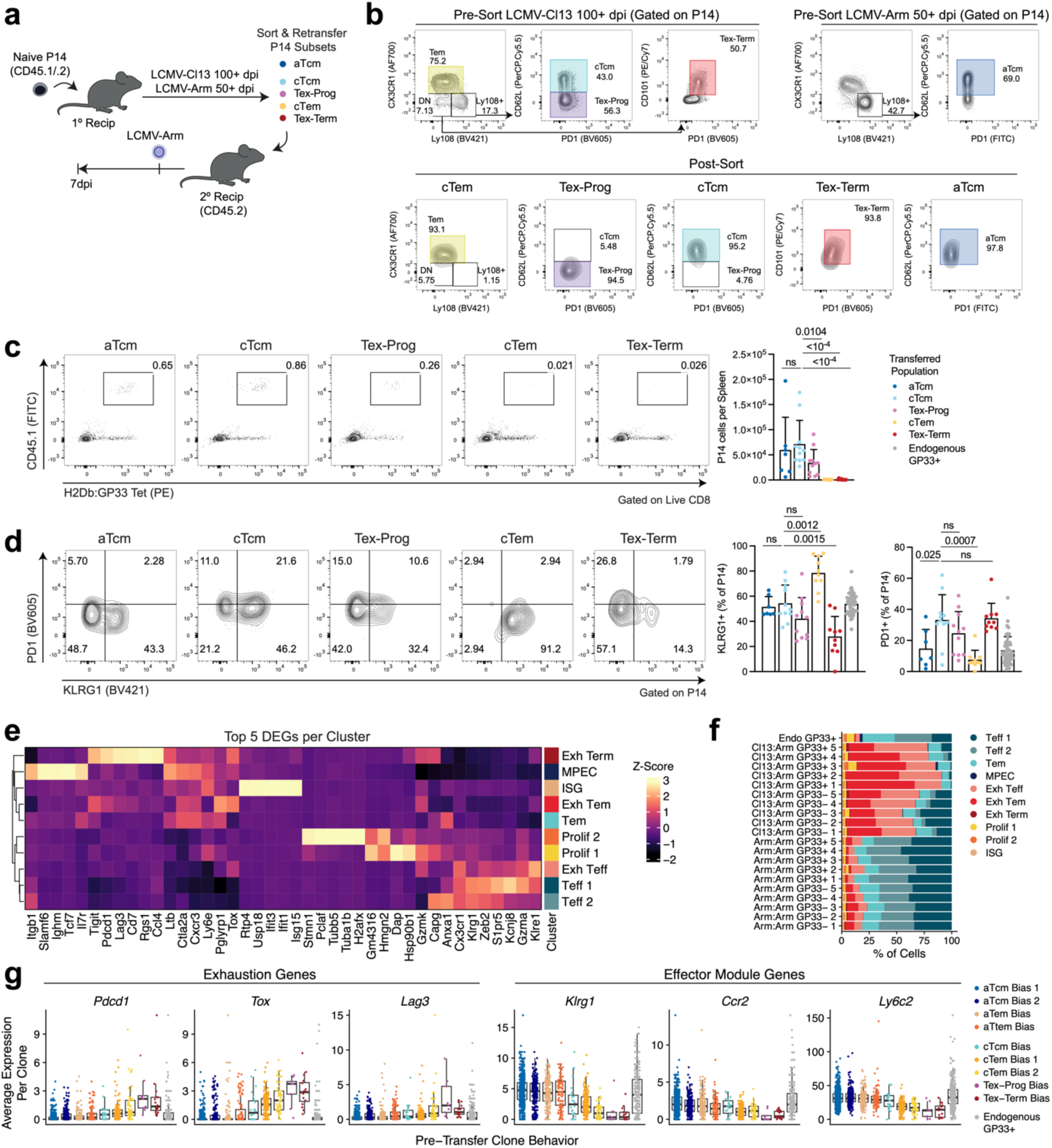
Functional recall response of chronic Tcm. (a-d) CD45.1^+^ CD45.2^+^ P14 subsets were isolated from LCMV-Arm or LCMV-Cl13 immune chimeras and 2×10^3^ cells were transferred into congenic hosts and challenged with LCMV-Arm. (b) Sorting scheme for P14 subsets and post-sort purity. (c) Flow cytometry quantifying recovery of P14s in secondary hosts 7dpi. (d) PD-1 and KLRG1 expression among P14 cells 7dpi. (a-d) Data are pooled from two independent experiments. Error bars show SD. n=7 aTcm, n=11 cTcm & Tex-Prog, n=10 cTem & Tex-Term. *P* values calculated by Mann-Whitney U Test. Data shown as mean+/−SD. ns=*P*>0.05. (e-g) scRNA/TCR-seq at 8 dpi from polyclonal Arm:Arm and Cl13:Arm rechallenge. (e) Top five differentially expressed genes (DEGs) per cluster in scRNA-seq of cells 8 dpi. (f) Cell frequencies from individual mice in cell clusters from scRNA-seq. (g) Expression of indicated genes from Cl13:Arm and Arm:Arm clones grouped by pre-transfer clone behaviors. Boxplots show Median, IQR, and 1.5xIQR.

**Extended Data Figure8:**
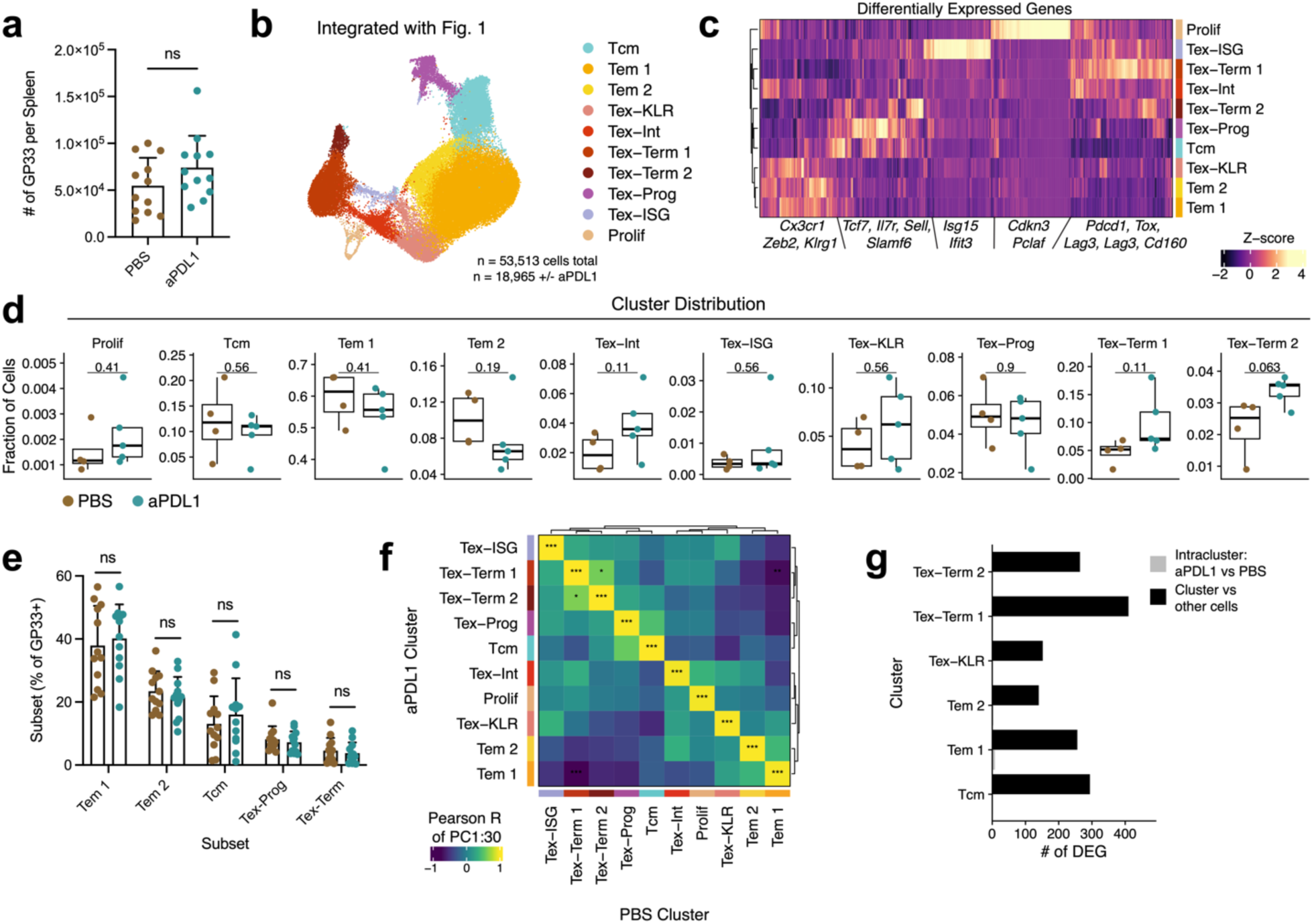
⍺PD-L1 treatment does not impact cell states of chronic memory cells. (a-g) Mice were infected with LCMV-Cl13 and treated with ⍺PD-L1 from 22 to 34 dpi. Flow cytometry and scRNA/TCR-seq of GP33^+^ cells were performed at 100+ dpi. (a) GP33^+^ cell number at per spleen 100+ dpi. (b) UMAP of scRNA-seq profiles GP33^+^ cells from PBS and ⍺PD-L1 treated mice integrated with data from Fig. 1 colored by cluster. (c) Differentially expressed genes (DEGs) from cell clusters. (d) Cluster proportion from individual mice in each group. Boxplots show Median, IQR, and 1.5xIQR. (e) Cell subset frequencies of GP33^+^ cells by flow cytometry. (f) Pearson correlation of PCs 1-30 of cells in each cluster between groups (Bonferonni corrected *P* values). (g) Number of DEGs between cells within the same cluster from PBS vs ⍺PD-L1 groups or between cells in cluster and all other cells. (a,e) n=12 mice/group. Data are pooled from two experiments. Bars show mean +/−SD. (b,c,d,f,g) n=4 mice PBS, n=5 mice ⍺PD-L1. (a-g) Unless otherwise noted, *P* values calculated by Mann-Whitney U Test. *=*P*<0.05, **=*P*<0.01, ***=*P*<0.001,ns=*P*>0.05.

## Methods

### Animal studies

All mouse studies were carried out in specific pathogen free facilities at Stanford School of Medicine in accordance with protocols approved by the Stanford Institutional Animal Care and Use Committee (protocol number: 33814). All mice used were female mice on the C57BL/6J background. Wild-type (JAX000064) and CD45.1^+^ (JAX002014) and TCRα^−/−^ (JAX002116) were purchased from Jackson Laboratories (Bar Harbor, ME). P14 TCR-transgenic mice were a gift from the Egawa Lab (Washington University, St. Louis) and crossed with C57BL/6J or CD45.1^+^ JAXBoy (JAX033076) mice for use in studies.

### LCMV infections

Mice were generally infected at 7-12 weeks of age. Mice were infected with LCMV-Cl13 by intravenous retro-orbital (r.o.) injection with 2×10^6^ plaque forming units (pfu) or LCMV-Arm by intraperitoneal (i.p.) injection with 2×10^5^ pfu. The same dose of LCMV-Arm was used for infection of TCRα^−/−^ mice. After 100 dpi LCMV-Cl13 infected mice were considered to have peripheral clearance of virus, however on occasion, viral titer remained high at this timepoint (**Extended Data Fig. 1a**), which was concordant with high (>75) % PD-1^+^ among GP33^+^ cells; these mice were excluded from experiments.

LCMV viral burden was quantified by qPCR as previously described^60^. Briefly, 250-500ng RNA extracted from Trizol (Invitrogen 15596-018) or spin columns (Zymo R1055) was used for cDNA synthesis (Quanta Bio 95048-100). qPCR was then performed with SYBR (BioRad 172-5275) and primers against LCMV-GP (fwd: cattcacctggactttgtcagactc, rev: gcaactgctgtgttcccgaaac) and quantified with a standard curve of linearized LCMV-GP containing plasmid (a gift from Miguel Sena-Esteves; Addgene 15793; ref^61^). To control for variability between experiments, LCMV viral burden data in Fig. 4l are presented as relative change from no transfer control.

### *α*PD-L1 treatment

Treatments were performed by i.p. injection of 200ug αPD-L1 monoclonal antibody (BioXCell clone B7-H1) every 2-3 days from 22-34 dpi. Control mice were i.p. injected with PBS on the same schedule.

### Cell isolation, flow cytometry, and cell sorting

Spleens were dissociated by mashing through 70-100μm filters into FACS buffer (PBS + 2% FBS + 1mM EDTA). Peripheral blood was collected r.o. with heparinized capillary tubes on anesthetized mice and blood was resuspended in FACS buffer. Cells were then either treated with ammonium chloride-potassium bicarbonate for red blood cell lysis or CD8^+^ T cells were enriched by magnetic selection (StemCell 19853). Single cell suspensions were stained with GP33:H2D^b^ Tetramer (MBL) for 30 minutes in FACS buffer at room temperature followed by Live/Dead dye and surface antibody staining in FACS buffer for 20-30 minutes on ice. FC receptor blocking was performed prior to antibody staining either in Tetramer staining mix or during CD8^+^ enrichment. For transcription factor staining, cells were fixed for 20 minutes in fixation / permeabilization buffer and stained for 1-2 hours in permeabilization buffer (eBioscience 00-5523-00). Prior to acquisition, stained cells were resuspended in FACS buffer and acquired on BD instruments (FACSAria III, Symphony, LSR-II). Sorting was performed with a BD FACSAria III into RPMI + 10-50% FBS at 4°C. Purity was routinely confirmed ≥90%. Flow cytometry analysis was performed in FlowJo (BD), and counting beads (Invitrogen C36995) were used to quantify absolute cell numbers. Antibodies and other reagents for flow cytometry used are listed in **Supplementary Table 8**.

### Adoptive T cell transfer

For all studies involving P14, naive CD8^+^ T cells were isolated from spleen and lymph nodes of P14 TCR-transgenic mice by magnetic selection as described above with the addition of 3ug/mL biotinylated αCD44 (Biolegend, clone IM7). 2×10^3^-4×10^3^ naive P14s were transferred per mouse. For other studies, CD8^+^ enriched spleens or sorted T cell subsets were used. For all experiments, cells were resuspended in PBS and injected into recipient mice by r.o. Injection. In experiments with subsequent infection, mice were infected the following day with LCMV. For fate mapping into LCMV-free mice, P14 cells were first enriched by magnetic selection of APC-CD45.1^+^ labeled cells (Miltenyi Biotec 130-090-855), followed by sorting, and transfer into LCMV-immune mice infected with LCMV-Arm >3 weeks prior were used to prevent carryover of virus^27,28^.

### Ex vivo quantification of cytokine secretion

1×10^3^ sorted cells from GP33^+^ subsets of T cells were resuspended in RPMI + 10% FBS. 50ng/mL PMA and 1ug/mL ionomycin (Biolegend 423302) was added to wells to a final volume of 80uL. Cells were stimulated at 37°C for 18 hours. 3-5 replicates were plated for each sorted population. Supernatant cytokine concentrations were assayed by cytokine bead array in duplicate (Biolegend 741050). Cytokine concentrations were fit to the standard curve in Qognit LEGENDplex software and differential cytokine production was calculated by ‘DESeq2’ with cTcm as a control group.

### scRNA/TCR-seq library preparation, data processing, and analysis

Prior to tetramer and antibody staining, CD8^+^ enriched spleens were stained with MULTI-seq lipid modified oligos (gift from Gartner Lab, UCSF) and barcodes as previously described^62^, except with a higher concentration of all reagents (µM) to accommodate increased cell number. Sorted CD8^+^ T cells were then pooled and up to 42,000 cells per well were loaded into a Chromium Controller and encapsulated in GEMs with 5’ v2 chemistry (10X Genomics 1000263). To improve integration, we included landmark cells in each well (Naive for Fig. 1 and endogenous GP33^+^ cells for Fig. 4) with a unique MULTI-seq barcode. 5’ scRNA, scTCR, and MULTI-seq libraries were prepared per manufacturer’s instructions or as previously described^62^. Indexed libraries were quantified using a Qubit and Agilent Bioanalyzer and pooled per manufacturer’s instructions. Pooled libraries were sequenced on a NovaSeqX (Illumina) instrument with the following parameters: 28bp Read 1, 10bp Index 1, 10bp Index 2, 90bp Read2.

scRNA-seq data was processed using CellRanger Count (v3.1.0; 10X genomics) aligned to the mm10 genome. scTCR-seq data was processed with CellRanger VDJ (v6.0.0; 10X genomics) and the ‘cellranger-vdj-GRCm38-alts-ensembl-3.1.0’ reference. Additional processing and analysis was performed with ‘Seurat’ (v4.3.0). We filtered cells by ‘nFeature_RNÀ (within 2 SD of mean and >1000), ‘percent.mt’ (< 95th percentile of dataset and < 4%), and biologically impossible TCR chain usage. After filtering, we utilized ‘deMULTIplex2’ to demultiplex samples and filter out doublets. We then performed ‘NormalizeDatà and ‘FindVariableFeatures’ (n=2000) on each 10X sample. To integrate data from multiple 10X samples, we implemented the reciprocal PCA pipeline. Integration features were found with ‘SelectIntegrationFeatures’, and TCR, cell cycle, mitochondrial, histone, and ribosomal genes were removed for all downstream analysis. Data was scaled and PCA was performed according to the integration features, and we ran ‘FindIntegrationAnchors’ and ‘IntegrateDatà to generate a single object. Seurat objects were then clustered and dimensionality reduction was performed by the standard pipeline. Briefly, we ran ‘ScaleDatà, ‘Run PCÀ, ‘FindNeighbors’, and ‘FindClusters’. For all datasets, we found clusters of low quality cells identified by high ‘percent.mt’ and high or low ‘nFeature_RNÀ or Naive cells based on gene expression (high Tcf7 and Sell; low Cd44 and Cxcr3) and/or clustered with intentionally sorted Naive cells. We removed unwanted cells and iteratively repeated clustering and dimensionality reduction. Final clustering and dimensionality reduction parameters: Fig. 1, PC1:12, Resolution=0.4; Fig. 4, PC 1:10, Resolution=0.4; Fig. 5, PC1:15, Resolution=0.34.

To identify differentially expressed genes between clusters, we utilized ‘FindAllMarkers’ (Wilcox Test, LogFC cutoff=0.25) on variable features with TCR, cell cycle, mitochondrial, histone, and ribosomal genes removed. To evaluate expression of exhaustion-related genes, we used ‘AddModuleScorè to create a gene module of the common exhaustion module from ref^11^ (**Supplementary Table 7**). To evaluate pairwise correlations of gene expression within groups and clusters, we selected cells from the most functionally important subsets of cells from each group, and found the median of each PC 1-30. Pearson correlation was calculated on this matrix to find correlations between groups, and we applied Bonferonni correction to *P* values. To find DEGs between aTem or cTem and other cells we grouped cells by cell type (ie Tcm, Tem, Tex-Term, Tex-Int, Tex-Prog, Tex-KLR) and group. We used ‘FindMarkers’ (Wilcox Test, Log_2_FC cutoff=0.25, *P*<0.01) on variable features with TCR, cell cycle, mitochondrial, histone, and ribosomal genes removed.

### Bulk TCR-seq library preparation and data processing

TCR-seq was performed by Adaptive Biotechnologies and iRepertoire. For genomic DNA based TCR-seq, samples were sorted, media was removed and cells were snap frozen. Immunosequencing TCR-seq was performed by Adaptive at the survey depth. For RNA based TCR-seq, cells were sorted into RNA lysis buffer (Zymo R1060) or DNA/RNA shield (Zymo R1100) and snap frozen. TCR-seq was performed by iRepertoire via their RepSeq+ Platform. Different sequencing methods were selected on a per-experiment basis based on trade-off of precision of absolute counts (better in genomic DNA-based), and detection of rare clones (better in RNA-based).

For both TCR-seq platforms, TCR sequence by count files were imported into R (v4.3.1) for analysis. For RNA-based TCR-seq, replicate libraries were amplified using each sample, so only TCR sequences detected between replicates were analyzed. For each TCR, we used the total number of cells in a subset and the clone’s frequency in each subset to calculate the total clone size and each clone’s phenotype distribution.

### ATAC-seq library prep, data-processing, and analysis

For ATAC-seq, GP33+ T cells were subset sorted from mice at different timepoints. Pre-sort, cells were combined into biologically distinct pools from multiple mice, and ATAC-seq was performed on these biological replicates. Tex-Term cells were also sorted from mice with cleared LCMV-Cl13, however, cell numbers were too low for reliable analysis.

Nuclei were isolated from 2,000-20,000 sorted T cells by resuspension in 50µL of Lysis Buffer (10 mM Tris-HCl pH 7.4, 10 mM NaCL, 3 mM MgCl_2_, 0.1% Tween-20, 0.1% IGEPAL, 0.01% Digitonin, 1% BSA) and spinning at 4°C 600rcf for 10 minutes. Nuclei were resuspended in 25uL Transposase master mix (2µL enzyme + 12.5µL 2X TD buffer + 10.5µL H_2_O; Illumina 20034198) and incubated at 37°C for 30 minutes with agitation at 400rpm. Tagmented DNA was isolated using MinElute PCR Purification Kit (Qiagen 28004) and amplified with Illumina adapters (ApexBio K1058) using Phusion PCR master Mix (New England Biolabs M0531S). Indexed libraries were size selected using SPRI (Beckman Coulter B23318) and quantified by Agilent Bioanalyzer, and pooled at equimolar ratios. Libraries were sequenced on a NovaSeqX (Illumina) instrument with 150bp paired end reads (1×10^8^ per library).

ATAC-seq libraries were analyzed using ‘PEPATAC’^63^ with default settings. Briefly, adapter sequences were trimmed with ‘--trimmer {skewer}’ followed by elimination of mitochondrial reads and reads associated with repetitive genomic regions. The remaining reads were aligned to the mm10 reference genome using Bowtie2. SAMtools was used to extract uniquely mapped reads and duplicate reads were removed using Picard; the resulting BAM files were used for downstream processing. OCR calling for individual samples was performed with MACS2, and non-overlapping 500-bp OCRs (also known as peaks) were generated. The OCRs by sample count matrix was constructed using default settings of the ‘run_atac’ function (ChrAccR). The matrix was variance-stabilized using the ‘vst’ function (DESeq2). Using the top 10,000 most variable OCRs, PC analysis plots were generated with ‘plotPCÀ (DESeq2) and samples were clustered by k-means clustering. Heatmap visualization was performed on the same set of 10,000 OCRs and the OCRs were clustered based on k-means clustering. To assess the number of differentially accessible OCRs between cTcm or Tex-Term and other cells, DESeq2 was used with a Log_2_FC cutoff of 1 and FDR cutoff of 0.01. For OCRs annotation to the nearest gene, the function ‘annotatePeak’ (ChIPseeker) was used. To find overlapping OCRs with data from ref^28^, we generated ‘GRanges’ objects from .bed files of DARs in each cluster from their figure 5c provided by authors, and found overlapping OCRs with our dataset using ‘findOverlaps’ (GRanges). Genome browser tracks were generated by aggregating data from replicate samples with ‘WiggleTools’ and plotting using ‘trackplot’ and ‘ggplot2’.

### Clone behavior analysis

To determine clone behavior of single GP33^+^ clones in scRNA/TCR-seq data, clones with a single *Tcra* and single *Tcrb* chain with >1 (Fig. 2a**, Extended Data Fig. 3a, Extended Data Fig. 4a,b**) or >2 (Fig. 5d) cells were isolated. For determination of clone behaviors in GP33^+^ bulk TCR-seq data (Fig 2b), we filtered expanded GP33^+^ clones with >4 cells sorted. For both data modalities, we clustered clones with k-means clustering based on frequency of the clone in each phenotype to determine clonal behaviors.

To determine if clones were significantly skewed towards any given phenotype versus the total population of observed cells (Fig. 2d) we performed a hypergeometric test as established in ref^64^. Specifically, we calculated a one-sided hypergeometric test of each clone’s frequency of the clone’s most frequent phenotype (n) compared to the total number of cells in the subset (N). We adjusted *P* values via Hochberg correction.

### Clone tracing between Tex and memory subsets

To determine if cTcm and other clone behaviors originated from clones present during chronic infection, we compared repertoires of GP33^+^ clones at 100+ dpi to GP33^+^ sorted (89.3 % PD-1^+^; **Extended Data Fig. 3d**) blood T cells at 28 dpi. To determine if clones from different behaviors are differentially detected at 28 dpi, we performed a binomial logistic regression with ‘glm’ and predicted if a clone could be detected in blood (n≥1 cell) based on the independent variable of 100+ dpi clone behavior.

To assess the clonal origins of cTcm and other clone behaviors in the exhausted pool (Fig. 2f**-i**), we analyzed subset sorted T cells from 21 and 100+ dpi. To expand breadth of response, we analyzed PD-1^+^ clones at 21 dpi and considered all clones detected in 21 dpi PD-1^+^ subsets as LCMV-specific. We restricted analysis to only clones >5 calculated sorted cells at both 21 dpi and 100+ dpi. To assess similarities between subsets, performed Pearson correlation on a matrix of individual clones by subset frequencies, adjusted *P* values with Bonferroni correction, and plotted correlations between subsets at different timepoints. To determine how clone behaviors at 21 dpi gave rise to different subsets at 100+ dpi, we clustered the shared clones separately based on clonal subset distribution at each timepoint. For the 100+ dpi timepoint, to resolve Tex-Prog and Tex-Term biased clones, we over-clustered (k=7) and combined two cTem 1 biased clusters into the same behaviors. To determine which clones preferentially formed cTcm biased clones, we performed logistic regression as described above with cTcm bias as the dependent variable and 21 dpi clone behavior as the independent variable.

To evaluate if blood sampling gave a representative view of clonal behavior in the spleen, we sampled PD-1^+^ Tex subsets from peripheral blood (collected via terminal cardiac puncture) and spleen of the same mouse (**Extended Data Fig. d-h**). We analyzed clone size of all clones and filtered on clones > 10 sorted cells in both organs to assess phenotype distribution between organs.

### Clonal tracing during polyclonal CD8^+^ rechallenge

To determine the clonal response of Arm:Arm and Cl13:Arm cells, we enriched polyclonal CD8^+^ T cells from three donor mice per group. Approximately half of cells were split into five recipient mice (2×10^5^ cells per mouse) and rechallenged, and the remaining cells were subset sorted for TCR-seq.

The first aim of this experiment was to confirm that rechallenge responses were clonally biased as has been shown in human memory T cells^41,43^. We isolated clones with a single *Tcra* and single *Tcrb* sequence and performed Pearson correlation of relative clone size (frequency of total cells) for each clone. To determine statistical significance, real clones were compared to a shuffled dataset where clone size in each host was shuffled and compared all correlations by Mann-Whitney. To compare gene expression profiles of clonally related cells in different hosts, we implemented PC correlation, which has been used to assess clonal similarity of gene expression in clonally barcoded hematopoietic stem cells^65^. First, we isolated single *Tcra* and single *Tcrb* clones with >10 cells and calculated cell-cell correlation based on the cell by PC correlation matrix for the PCs 1-30 of gene expression data. We then compared correlation between clonal sisters in the same hosts and other hosts. To determine significance of this correlation, we generated random clones by shuffling the clone id and compared this value to real clones.

To assess expansion of individual clones, we filtered clones single *Tcra* and single *Tcrb* clones which were detected at both timepoints. We performed no further filtering for clonesize to ensure representation of small clones. To assess expansion, we calculated the Log_2_FC of post total number of cells per clone at 8 dpi (clone frequency*total number of cells recovered) versus pre-rechallenge (clone frequency*total number of cells transferred). To determine which clonal behaviors of cells expanded best upon rechallenge, we performed k-means clustering of phenotype proportion data of Arm:Arm and Cl13:Arm clones pre-rechallenge. Expansion of clones in different pre-rechallenge clone behaviors were compared by Mann-Whitney U test.

To determine the gene expression profiles of cells from different pre-rechallenge clone behaviors, we analyzed the same set of clones, and further filtered on clones with >4 cells in the 8 dpi dataset. We calculated the probability of clones in a given pre-rechallenge behavior to derive clones predominantly in the exhaustion-associated clusters (>50% of clone’s 8 dpi progeny in ‘Exh Teff’, ‘Exh Tem’, or ‘Exh Term’ clusters) by logistic regression as described above (independent variable = pre-rechallenge clone behavior, dependent variable = >50% in exhaustion-associated clusters). To determine how levels of effector molecules differed among individual Cl13:Arm clones, we added an effector module based on the top DEGs from the effector cell cluster (top 50 DEGs, filtered for presence in variable features in our dataset) from CD8^+^ T cells responding to LCMV-Arm infection (ref^11^; **Supplementary Table 7**) using ‘AddModuleScorè (Seurat). To determine if individual clones had a significantly different effector module score from the total distribution of Cl13:Arm cells, we performed a two-sided Mann-Whitney test with Benjamini & Hochberg correction of individual clones versus the total distribution of Cl13:Arm clones as established in ref^41^. We then quantified the number of clones with significantly (*P*<0.05) higher or lower effector module score than the total distribution of Cl13:Arm cells.

### Analysis of clonal behavior with αPD-L1 treatment

For clustering of clones in scRNA/TCR-seq after ICB (Fig. 5d), we over-clustered clones (k=8) and combined multiple cTem 1 biased behaviors to resolve Tex-Prog and Tex-Term biased clones. To assess clonal responses to αPD-L1 treatment, we analyzed subset sorted T cells from 21 dpi, 35 dpi, and 100+ dpi (Fig. 5e**-j**). Like with above, we assessed total T cells as to not restrict analysis to a single antigen specificity, and analyzed only cells that were detected in PD-1^+^ subsets at either 21 or 35 dpi. For all analysis, we filtered clones on clones ≥10 sorted cells. We first assessed clonal behaviors of exhaustion derived clones which we traced to the late timepoint. To determine longitudinal trajectories of clones, we based analysis on the pipeline outlined in ref^48^, which is designed to identify common behaviors of clones at multiple timepoints. We modified this pipeline to take into account both total clone size and phenotype proportion. We created a matrix of clones by phenotype frequency (clonotype subset count/total cell count at timepoint) at all timepoints. To identify clones with common clonal trajectories over time, we calculated Pearson correlation of each set of clones and performed hierarchical clustering of clones based on euclidean distance of the correlation matrix. To link longitudinal trajectories to memory cell state, we utilized a hypergeometric test to determine if clones were biased towards a phenotypic subset. We calculated a one-sided hypergeometric test of each clone’s frequency in each subset (n) compared to the total number of cells in the subset (N). We adjusted *P* values via Hochberg correction, and clonal bias was determined by a clones’ lowest significant *P* value (*P*<0.05). To identify clone trajectory enrichment the αPD-L1 group, logistic regression was performed as described above with group as the dependent variable and clone behavior as the independent variable.

### Statistical analysis

All statistical analyses were performed in R (v4.3.1) and GraphPad Prism (v10). Statistical tests for each experiment are noted in the above methods and/or figure legends. Unless noted, all statistical tests were two-sided. No statistical testing was performed to pre-determine sample size.

## Supplementary Tables

Supplementary Table 1 - Cluster DEGs of scRNA/TCR-seq from GP33^+^ atlas

Supplementary Table 2 - DEGs between cTcm or cTem and other subsets

Supplementary Table 3 - 10,000 most variable OCRs from ATAC-seq and Z-score accessibility per sample

Supplementary Table 4 - DARs between Tex-Term or cTcm and other populations

Supplementary Table 5 - Cluster DEGs from scRNA/TCR-seq post rechallenge

Supplementary Table 6 - Cluster DEGs from scRNA/TCR-seq after aPDL1 treatment

Supplementary Table 7 - Gene module scores

Supplementary Table 8 - Flow cytometry reagents

Supplementary Table 9 - Source data

## Data Availability

Raw sequencing data from this study are available on NCBI GEO at accession numbers GSE285411 (scRNA/TCR-seq atlas), GSE285412 (scRNA/TCR-seq αPD-L1), GSE285414 (scRNA/TCR-seq post-rechallenge) and GSE285308 (ATAC-seq). Bulk TCR-seq data are available on Zenodo (doi.org/10.5281/zenodo.14648171). Source data for other experiments are provided in **Supplementary Table 9.**

## Code Availability

No custom algorithms were developed for this study. Code and processed data needed to generate figures from this study are publicly available online at GitHub at: github.com/colinraposo/ChronicCentralMemory.

## Acknowledgements

This work is supported by the National Institutes of Health (NIH) U01CA260852 (A.T.S.), a Career Award for Medical Scientists from the Burroughs Wellcome Fund (A.T.S.), a Lloyd J. Old STAR Award from the Cancer Research Institute (A.T.S.), the Parker Institute for Cancer Immunotherapy (A.T.S.), and a Pew-Stewart Scholars for Cancer Research Award (A.T.S.). K.J.H-G. is supported by a Stanford School of Medicine Propel Postdoctoral Scholarship and Cancer Research Institute Irvington Postdoctoral Fellowship. C.J.R. was funded by an NIH training grant through the Stanford Immunology Program (5T32AI007290-38). Sequencing was performed at the UCSF CAT, supported by UCSF PBBR, RRP IMIA, and NIH 1S10OD028511-01 grants. Flow cytometry was partially performed with instruments in the Stanford Shared FACS Facility (RRID: SCR_017788) on instruments obtained by NIH grants (S10RR027431-01 and 1S10OD026831-01).

## Author Contributions

C.J.R., K.J.H-G, and A.T.S. conceptualized the study. C.J.R., K.J.H-G, and A.T.S. wrote and edited the manuscript with input from all authors. C.J.R., P.K.Y., S.M., and K.J.H-G. performed experiments. C.J.R., A.Y.C., and K.J.H-G. performed data analysis. K.J.H-G. and A.T.S. guided experiments and data analysis.

## Declaration of Interests

A.T.S. is a founder of Immunai, Cartography Biosciences, Santa Ana Bio, and Prox Biosciences, an advisor to 10x Genomics and Wing Venture Capital, and receives research funding from Astellas and Northpond Ventures. All other authors declare that they have no competing interests.

## Notes

https://github.com/colinraposo/ChronicCentralMemory

